# Cell types and clonal relations in the mouse brain revealed by single-cell and spatial transcriptomics

**DOI:** 10.1101/2021.08.31.458418

**Authors:** Michael Ratz, Leonie von Berlin, Ludvig Larsson, Marcel Martin, Jakub Orzechowski Westholm, Gioele La Manno, Joakim Lundeberg, Jonas Frisén

## Abstract

The mammalian brain contains a large number of specialized cells that develop from a thin sheet of neuroepithelial progenitor cells^1,2^. Recently, high throughput single-cell technologies have been used to define the molecular diversity of hundreds of cell types in the nervous system^3,4^. However, the lineage relationships between mature brain cells and progenitor cells are not well understood, because transcriptomic studies do not allow insights into clonal relationships and classical fate-mapping techniques are not scalable^5,6^. Here we show *in vivo* barcoding of early progenitor cells that enables simultaneous profiling of cell phenotypes and clonal relations in the mouse brain using single-cell and spatial transcriptomics. We reconstructed thousands of clones to uncover the existence of fate-restricted progenitor cells in the mouse hippocampal neuroepithelium and show that microglia are derived from few primitive myeloid precursors that massively expand to generate widely dispersed progeny. By coupling spatial transcriptomics with clonal barcoding, we disentangle migration patterns of clonally related cells in densely labelled tissue sections. Compared to classical fate mapping, our approach enables high-throughput dense reconstruction of cell phenotypes and clonal relations at the single-cell and tissue level in individual animals and provides an integrated approach for understanding tissue architecture.

## Main

The mammalian brain contains a vast number of specialized cell types organized into networks that perform computations and enable complex cognitive processes. Ever since the discovery of various cell types in the nervous system, our understanding of cell diversity in the mammalian brain has been increasingly refined. Modern high-throughput technologies such as single-cell RNA sequencing (scRNA-seq) revealed the existence of hundreds of molecularly distinct cell types including diverse types of neurons, astrocytes, oligodendrocytes and microglia across the entire mouse and human nervous system^3,7-11^. Yet, our molecular understanding of the developmental origins of cell diversity remains limited and a systematic analysis of lineage relationships between brain cells is lacking due to the low throughput of classical fate mapping techniques^5,6^. Advanced molecular tools based on genome editing or genetic barcoding have been used to record cell lineages^12-17^, and combined with scRNA-seq to generate fate maps in cultivated cells^18,19^, zebrafish^20-23^ and mice^17,19,24,25^. However, these technologies are not readily employed to uniquely label a large number of neuroepithelial progenitor cells in the mouse brain *in vivo* and most approaches require tissue dissociation. Establishing a system that allows genetic barcoding at large scale with an *in situ* whole transcriptome readout is crucial for studies of the nervous system where function arises from both differential gene expression and circuit-specific anatomy^26-29^.

Here we describe TREX, a robust and high-throughput approach that allows for simultaneous clonal *tr*acking and *ex*pression profiling of mouse brain cells and Space-TREX, which utilizes spatial transcriptomics coupled to immunohistochemistry for tracking clonally related cells in their native environment in tissue sections. Both methods are based on the intraventricular injection of a diverse lentivirus library into the neural tube of the mouse embryo to deliver a compact, heritable and ubiquitously expressed genetic barcode into precursor cells. We applied TREX to study the lineage potential of early neuroepithelial cells and revealed the existence of fate-restricted progenitor cells as early as E9.5 for the murine hippocampus. We uncovered that microglia, the tissue-resident macrophages of the brain, are generated from a limited number of progenitor cells that undergo massive clonal expansion as well as widespread migration across multiple regions of the mouse telencephalon. Using Space-TREX, we demonstrate that migration patterns of progeny from brain progenitor cells can be investigated using spatial transcriptomics. Together, our findings demonstrate the utility of TREX and Space-TREX for high-throughput clonal tracing in the mouse brain and provide new tools and molecular insights to study brain development at the single-cell and tissue level. These techniques are easy to use with standard laboratory equipment and can be applied to any model organism to study the development of any tissue of interest.

## Results

### Unique and heritable labelling of progenitor cells with expressed barcodes

Here, we present TREX, a lentivirus-based strategy for barcoding of mouse brain progenitors *in vivo* followed by single-cell transcriptomics for tracking clonal relationships and profiling gene expression of thousands of brain cells in one assay (**Fig. 1a**). Our approach relies on a diverse lentivirus library containing random barcodes or cloneIDs with a length of 30 bp downstream of a nuclear-localized EGFP driven by a strong, ubiquitous EF1a promoter (**Extended Data Fig. 1a-b**). To ensure high library complexity required for unique labelling of progenitor cells, we used next generation sequencing of lentivirus preparations (**Extended Data Fig. 1c-f**). We found 1.57 × 10^6^ ± 0.12 × 10^6^ cloneIDs/µl (mean ± SD, n = 4 preparations) in a typical lentivirus preparation with a largely uniform representation (Gini index = 0.2) and high sequence diversity (Hamming distance = 22 ± 2.4; mean ± SD; n = 10,000 random samples).

**Fig. 1.**
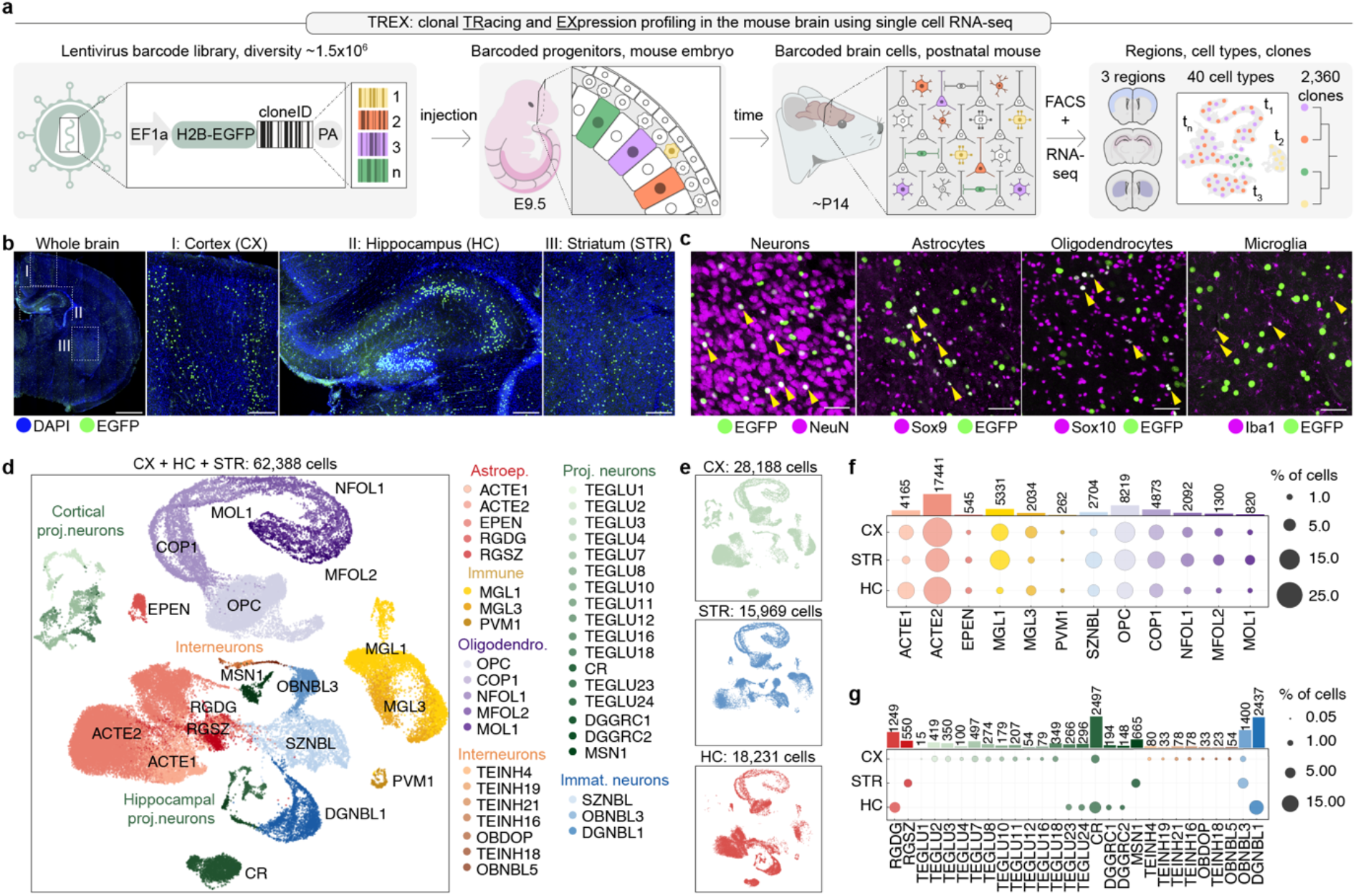
TREX enables simultaneous profiling of cell phenotype and clonality. **a**, Workflow for *in vivo* barcoding and profiling of brain cells. Microinjection of a highly diverse lentiviral barcode library into the ventricle of the developing mouse brain results in the expression of the barcode and EGFP in progenitor cells and their progeny. Each barcode serves as a unique cloneID that is inherited as progenitors differentiate into diverse brain cell types and single-cell RNA-seq is used to reveal the transcriptome and cloneID. We injected virus at embryonic day (E) 9.5 and isolated barcoded cells from three forebrain regions at postnatal day (P) 14 for RNA-seq which revealed 40 cell types and a total of 2,360 clones. **b**, Barcoding at E9.5 results in widespread and stable transgene expression in telencephalic regions including cortex (CX), hippocampus (HC) and striatum (STR) of the postnatal brain. Scale bars: whole brain, 1 mm; crop outs, 100 µm. **c**, Barcoding at E 9.5 results in stable transgene expression in all major cell types of the postnatal mouse brain. Scale bar: 50 µm. **d**, Visualization of identified cell classes using uniform manifold approximation and projection (UMAP). In total, 62,388 single cell transcriptomes were collected from three telencephalon regions and five brains (barcoded and non-injected controls) that were classified into 40 cell types. Capital black letters indicate a unique identifier for each cell type taken from www.mousebrain.org. Colours indicate six broader cell type classes: astroependymal (reds), immune (yellows), interneurons (oranges), projection neurons (greens), immature neurons (blues) and oligodendrocytes (purples). **e**, The same UMAP as in panel d split by regional origin of cells: cortex (n = 28,188 cells, top), striatum (n = 15,969 cells, middle) and hippocampus (n = 18,231 cells, bottom). **f, g**, Dot plots showing fractions of cells in each class per region for cell types found in all three regions (**f**) and for cell types unique to one or two regions (**g**). Bar plots show total numbers of cells for each cell type.

To label individual progenitor cells *in vivo*, we used *in utero* microinjection of lentivirus into the ventricular system of the mouse forebrain at embryonic day (E) 9.5 (**Extended Data Fig. 2a**). We typically injected about 0.6 µl of EGFP-cloneID virus corresponding to 0.94 × 10^6^ unique cloneIDs and quantified the number of barcoded cells two days after virus injection at E11.5. Analysis of single-cell suspensions from whole brains at E11.5 revealed labelling of 1.8 ± 0.25% of cells (mean ± SD, n = 3 brains, **Extended Data Fig. 2b**) corresponding to a total number of 41,000 ± 3,500 cells (mean ± SD, n = 3 brains) per E11.5 mouse brain (**Extended Data Fig. 2b, c**). Because the cell cycle length is around 12 h during this time of brain development^30^, we estimated that the initial number of labelled progenitor cells at E9.5 was around 2,600 and that 99.6% of cells were uniquely labelled with a cloneID (**Extended Data Fig. 2d**). However, even if more cells were to be barcoded, our lentivirus library is sufficiently complex to uniquely label 99% of a total of 6,310 cells.

**Fig. 2.**
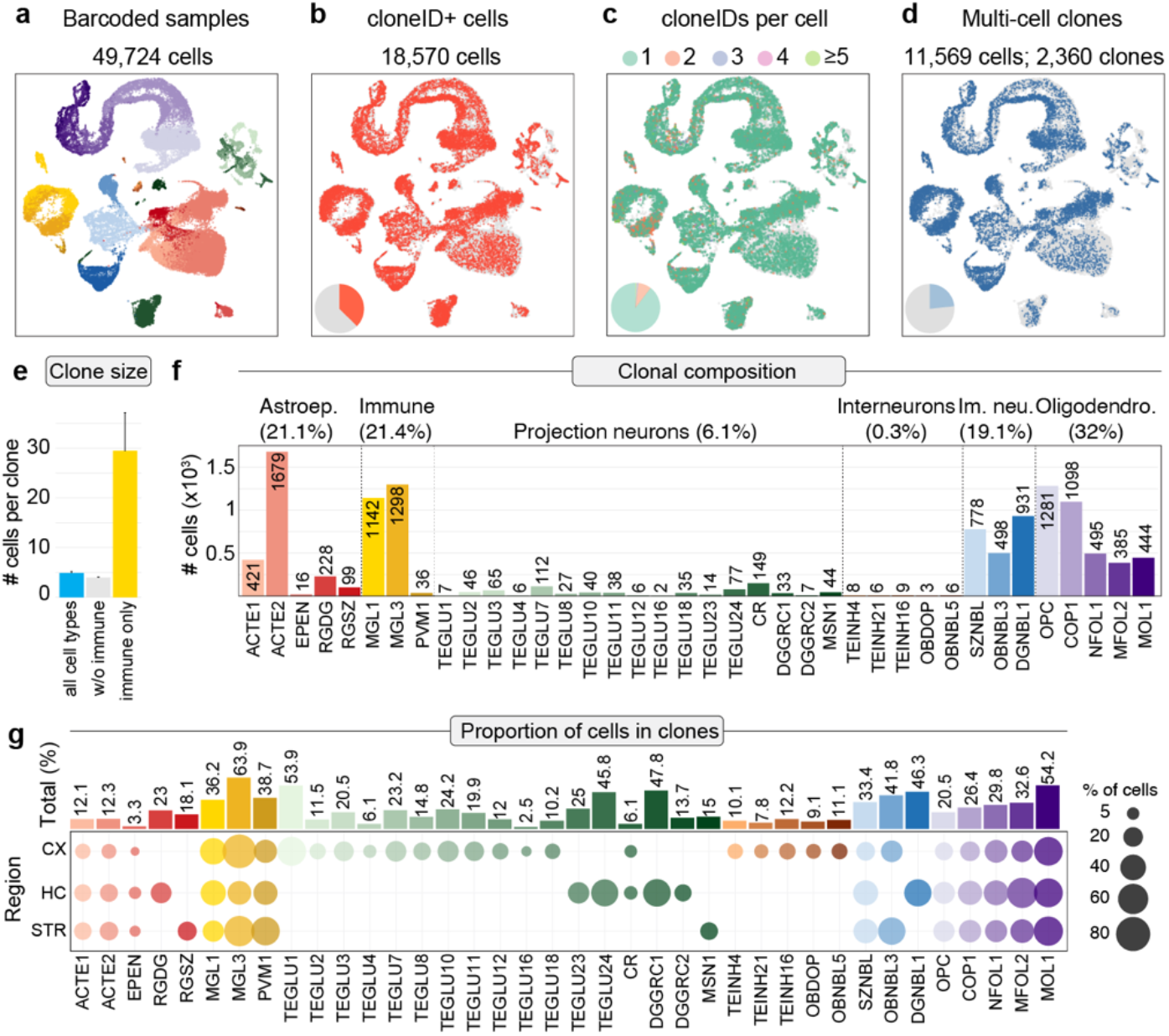
Clone reconstruction across cell types and regions in the mouse telencephalon. **a-d**, UMAP visualizations of all cells isolated from barcoded samples grouped by cell type (a), cloneID-expressing cells (b), number of cloneIDs per cell (c) and cells in multi-cell clones (d). **e**, Bar plots showing average clone sizes (number of cells per clone). The average clone size varies depending on cell type and is 4.9 ± 0.3 cells/clone when considering all cell types (left bar; n = 2,360 clones), 4 ± 0.1 cells/clone when considering only neuroectoderm-derived cells (middle bar, n = 2,276 clones) and 29.5 ± 7.6 cells/clone when considering only mesoderm-derived immune cells (right bar, n = 84 clones). Values correspond to mean ± SEM. **f**, Bar plots showing total number of cells in clones for each cell type. **g**, Dot plots and bar plots displaying the total proportion of cells in clones for each cell type (top) and per region (bottom).

Barcoded EGFP+ cells were mostly evenly distributed throughout the E11.5 neuroepithelium and included Sox2+ radial glia progenitors lining the ventricular zone as well as their Sox2-daughter cells (**Extended Data Fig. 2e-g**). Long-term EGFP-cloneID expression was maintained in the juvenile mouse brain and labelled cells were found in various regions such as the cortex, hippocampus and striatum (**Fig. 1b**). Antibody staining with markers against major cell types of the brain showed that neurons (Rbfox3/NeuN), astrocytes (Sox9), oligodendrocytes (Sox10) and microglia (Iba1) were labelled with a heritable cloneID (**Fig. 1c**). In conclusion, we present a highly diverse lentivirus library suitable for unique labelling of mouse brain progenitor cells with barcodes that display long-term expression in all major cell types of the postnatal brain.

### Single-cell profiling reveals the molecular identity of barcoded brain cells

To analyse the fate potential of barcoded progenitor cells and to determine the molecular identity of their daughter cells, we applied TREX to cells labelled at E9.5. We dissected brains from two-week-old mice and used fluorescence-activated cell sorting (FACS) to isolate all EGFP+ barcoded cells separately from cortex, striatum and hippocampus for scRNA-seq (**Extended data Fig. 3a-d**). We collected cells from these three regions per brain from five brains (four barcoded and one non-injected control) and used droplet microfluidics to reveal the transcriptome profiles of 65,160 cells. Graph-based clustering revealed five main clusters corresponding to astroependymal cells, immune cells, neurons, oligodendrocytes and vascular cells (**Extended data Fig. 3e**). Both control and barcoded samples showed a similar cell type composition with the exception of vascular cells which represented 16.6% of cells in the control dataset and less than 0.5% of barcoded cells (**Extended data Fig. 3f**). This can be attributed to the fact that blood vessels only begin to sprout into the ventrolateral brain at E9.5 (ref^31^), which results in a low number of cells that can be labelled at the time point of injection. We therefore removed the cluster of vascular cells from all datasets and kept a final of 62,388 single-cell profiles with a mean of 5,444 transcripts and 2,255 genes detected per cell (**Extended data Fig. 3g-l**).

**Fig. 3.**
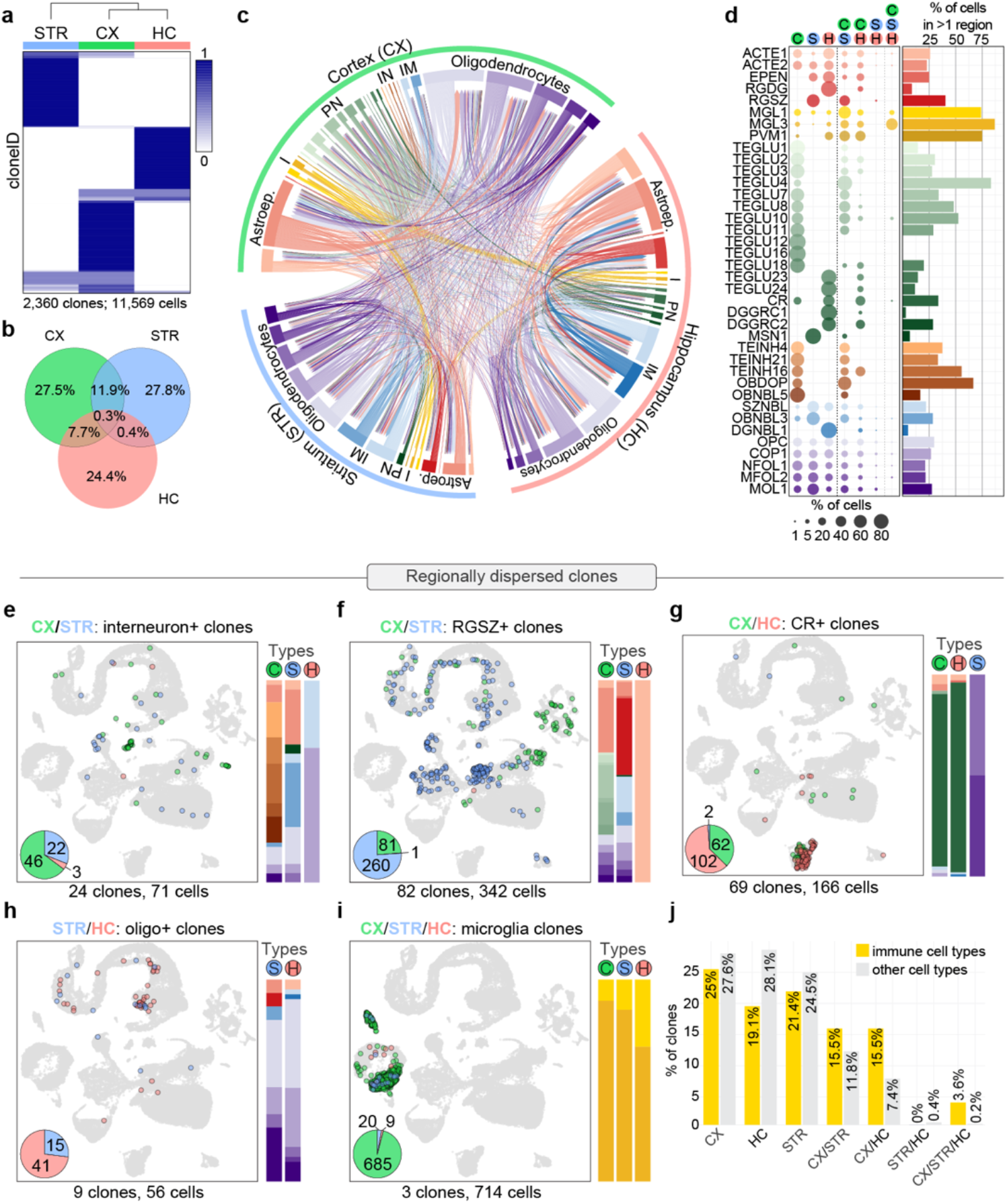
Regional distribution of clonally related cells across the telencephalon. **a, b**, Most cells belonging to the same clone were restricted to a single brain region. For each cloneID (rows), the proportions of cells within each region (columns) were calculated, scaled by row and coloured as shown in the figure (a). The fraction of cells per clone belonging to one region (CX, STR or HC), two regions (CX/STR, CX/HC or STR/HC) or all three forebrain regions (CX/STR/HC) was determined and displayed as Venn diagram (b). All 2,360 clones and a total of 11,569 cells were considered. CX, cortex (green); STR, striatum (blue); HC, hippocampus (red). **c**, *Circos* plot displaying shared cloneIDs between all cell classes (inner segments) across CX, STR and HC (outer segments). For each cell pair, the number of shared cloneIDs is indicated by the width of the link and the colour of each link represents cell type. **d**, Cell types associated with dispersed clones were identified by determining the cell type composition of clones spread across multiple telencephalon regions relative to the total number of cells in clones for each cell type. The bar plot summarizes the proportion of cells in clones spread across multiple regions. **e-i**, Examples of cell types in clones dispersed across multiple regions. Each example contains a UMAP visualization of cells, a pie chart displaying the number of cells per region and bar plots to illustrate the cell type composition of clones. Selected clones dispersed across the anatomical boundaries between cortex and striatum (e, f), cortex and hippocampus (g), striatum and hippocampus (h) and all three regions (i) are displayed. **j**, Clonally related immune cells (MGL1, MGL3, PVM1) disperse more frequently across telencephalon regions compared to neuroectoderm-derived cells. The fraction of clones containing clonally related cells found in one region (CTX, STR, HC), two regions (CX/STR, CX/HC, STR/HC) or three regions (CX/STR/HC) is shown.

Next, we performed subclustering for each major cell type from all brain regions and assigned each cell subclass a unique mnemonic identifier based on an existing mouse brain atlas^3^ (**Extended data Fig. 4**). In total, we found 40 molecularly defined cell classes with the largest diversity among neuronal cells which included 17 types of projection neurons, 7 types of GABAergic interneurons and 3 types of immature neuronal cells (**Fig. 1d**). As expected, clusters corresponding to glial cell types were less heterogenous and we found 5 subclasses of astroependymal cells, 5 subclasses of oligodendrocyte lineage cells and 3 subtypes of immune cells. From the three telencephalic regions sampled we collected the highest number of cells from cortex (n = 28,188 cells) followed by hippocampus (n = 18,231 cells) and striatum (n = 15,969 cells) (**Fig. 1e**). Certain cell types such as protoplasmic astrocytes (ACTE2, n = 17,441 cells), microglia (MGL1, n = 5,331 cells) or mature oligodendrocytes (MOL1, n = 820 cells) were found in similar proportions in each region (**Fig. 1f**). However, many cell types were specific for one region such as dentate gyrus radial glia-like cells in the hippocampus (RGDG, n = 1,249 cells), medium spiny neurons in the striatum (MSN1, n = 665 cells) and upper layer projection neurons in the cortex (TEGLU7, n = 497 cells) (**Fig. 1g**). Finally, we compared gene expression profiles and total cell type composition between barcoded and non-injected samples which indicated that lentivirus-mediated barcoding does not perturb cell physiology (**Extended data Fig. 5**). Together, these data show the utility of TREX for barcoding progenitor cells in the developing brain and profiling the identity of their barcoded progeny at a postnatal stage using single-cell RNA-seq.

### Barcode expression metrics across cell types

To specifically study barcoded cells, we removed cells from non-injected control samples from the full dataset (n = 62,388 cells) and focused on the 49,724 cells isolated from four mouse brains that were previously injected with an EF1a-EGFP-cloneID virus library (**Fig. 2a**). We detected EGFP transcripts in a total of 21,743 cells (43.7%) corresponding to 44.9% to 68.5% (57.4% ± 10.2%, mean ± SD, n = 4 brains) EGFP mRNA positive cells per brain (**Extended Data Fig. 6a, b**). The average number of EGFP transcripts per cell was highest in immune cells followed by intermediate levels in immature neurons, projection neurons, oligodendrocytes as well as interneurons and lowest expression levels in astroependymal cells (**Extended Data Fig. 6c**). The number of EGFP transcripts per cell class were correlated (r = 0.81) with Elongation Factor 1-Alpha 1 (Eef1a1) levels indicating that transgene expression under the synthetic EF1a promoter faithfully recapitulates endogenous Eef1a1 expression patterns albeit at lower levels (**Extended Data Fig. 6d**).

Using a custom pipeline, we extracted cloneIDs directly from single-cell transcriptome data as well as targeted amplicon libraries (**Extended Data Fig. 7a, b**). We captured a total of 21,433 cloneIDs in 18,570 cells (37.3%) corresponding to 24.2% to 50.9% (38.4% ± 11.8%, mean ± SD, n = 4 brains) of all sampled cells per brain (**Fig. 2b; Extended Data Fig. 7c, d**). We captured cloneIDs for most of the previously identified 40 cell types except for one very rare type of interneurons, TEINH18. The average number of cloneIDs per cell was similar when using scRNA-seq (1.15 cloneIDs per cell on average) or bulk DNA sequencing (1 cloneID per cell on average) of barcoded cells (**Extended Data Fig. 7e, f**) suggesting that cloneID capture is quantitative using single-cell transcriptomics.

While the vast majority of cloneID+ cells (89.6%) across all brains expressed only one cloneID (**Fig. 2c**), the proportion of such cells varied between brains and ranged from 78.6% to 95.8% with the remaining fraction of cells expressing multiple cloneIDs (**Extended Data Fig. 7g, h**). Based on the transduction rate of 1.8% (see above) and an idealized transduction model^32^, we expected that 99.08% of cells contain one cloneID and 0.92% of cells contain two or more cloneIDs (**Extended Data Fig. 7i**). Although we followed a rigorous strategy for doublet removal (see methods), we were concerned that cells with multiple cloneIDs could correspond to undetected doublets in our scRNA-seq data. Therefore, we injected multicolour lentiviruses into the embryonic forebrain and found that 3.4x more cells than expected contained multiple fluorophores (n = 1,301 cells) (**Extended Data Fig. 7j-l**). We found that the observed deviation from the theoretical cloneID copy number distribution per brain can be attributed to (1) variable copy numbers per cell type and (2) variable proportions of barcoded cell types in each brain (**Extended Data Fig. 8**). First, the total fraction of cells expressing multiple cloneIDs ranged from about 3% in white matter astrocytes (ACTE1) to 16.7% in some excitatory cortical projection neurons (TEGLU1, TEGLU10) and was highest in immune cell types: 17.9% of MGL1, 31.2% of PVM1 and 43.2% of MGL3 contained more than one cloneID. The correlation coefficients between the fraction of cells with multiple cloneIDs and cloneID expression levels were highly variable (−0.96 ≤ r ≤ 0.97) for each major cell type indicating that cloneID transcript levels can only partially explain the number of cloneIDs per cell type (**Extended Data Fig. 9**). Instead, increased transduction rates of local progenitor cells due to position and/or differential expression of receptors required for lentivirus entry^33^ might result in a higher occurrence of certain cell types harboring more than one cloneID sequence, even at low transduction rates. For example, the choroid plexus serves as an entry site for microglia progenitors to first enter the cerebrospinal fluid and then the brain at the ventricular surface^34,35^ and we observed “hot spots” with a higher density of EGFP+ barcoded cells in the ventral and dorsal regions of the E11.5 neuroepithelium (**Extended Data Fig. 2e**) corresponding to the developing choroid plexus–cerebrospinal fluid system^34^. Second, each brain contained different amounts of barcoded cells from certain cell types, e.g. brains 1 and 3 contained more microglia and less ACTE1 than the other brains leading to variable cloneID copy number distributions for each brain. In summary, all major brain cell types were represented among barcoded cells and most cells express a single cloneID with varying copy numbers per cell type.

### Clonal relationships across forebrain regions

We identified clonally related cells based on the Jaccard similarity of cloneIDs for each pair of cloneID containing cells^18^ and we defined clones as groups of two or more related cells. In total, we reconstructed 2,360 clones containing 11,569 cells (23.3 % of all cells; **Fig. 2d**) with an average size of 4.9 ± 0.3 cells per clone (mean ± SEM) and the number of clones per brain ranged from 201 to 1,106 (11.1% to 38.6% of all cells per brain; **Extended Data Fig. 10**). Interestingly, clones containing mesoderm-derived myeloid cells (microglia and perivascular macrophages) were about 7.4 times larger than those with neuroectoderm-derived cells and contained 29.5 ± 7.6 cells per clone (mean ± SEM, n = 84 clones, max = 534 cells per clone) compared to 4 ± 0.1 cells per clone (mean ± SEM, n = 2,276 clones, max = 116 cells per clone), respectively (**Fig. 2e**). This difference in clone size probably reflects the massive proliferation of brain macrophages required to colonize the entire CNS, after only a small number of precursors enters the brain before closure of the blood-brain barrier around E13, restricting access to immune cells that arise later in development^36^.

To estimate the potential error associated with clone reconstruction, we quantified how often cell types that arise from different progenitors shared the same cloneID. First, we found that clones containing cortical excitatory (n = 371 clones) or inhibitory neurons (n = 18 clones), which are known to have separate developmental origins^37^, never shared the same cloneID (**Extended Data Fig. 11a**). Second, among 84 clones containing 2,481 mesoderm-derived microglia or perivascular macrophages only 3 clones with a total of 453 cells shared a cloneID with 5 neuroectoderm-derived cells (**Extended Data Fig. 11b-d**) and we removed these cells from the respective clones. These data suggest a low error rate of about 0.2% (5 out of 2481 cells) that could be related to clone size and cell type since only large immune clones (3 clones, 71 to 219 cells per clone) contained neuroectoderm-derived cells or to non-unique cloneID labelling. Moreover, we found that cloneID removal from cell types that often express more than one cloneID such as mesoderm-derived cells are affected by “lumping” errors leading to less clones with a larger size and a higher number of incorrectly associated neuroectoderm-derived cells (**Extended Data Fig. 12**). This is expected since the co-expression of two or more distinct cloneIDs per cell leads to a higher combinatorial diversity^18^ thus reducing the error associated with clone reconstruction. Finally, there was a high correlation (r = 0.99) between barcode frequency and the number of cells with a barcode in distinct clones indicating that there is no preferential uptake of certain barcodes among progenitor cells leading to uniform library representation of labelled cells (**Extended Data Fig. 13**).

Cell types most often represented in clones were oligodendrocyte subtypes (3,703 cells, 32%) followed by immune cells (2,476 cells, 21.4%), astroependymal cells (2,443 cells, 21.1%), immature neuronal cells (2,207 cells, 19.1%), projection neuron types (708 cells, 6.1%) and interneurons (32 cells, 0.3%) (**Fig. 2f**). Except for one type of interneurons, TEINH19, we captured clonal information for all cell types that also contained a cloneID. Cell types containing the highest proportion of cells in clones were cortical and striatal microglia (MGL3) of which 77.5% and 60.6%, respectively, of all sampled MGL3 cells per region were represented in clones (**Fig. 2g**). Similarly, 52.5% to 55.1% of all mature oligodendrocytes (MOL1) were found in clones across all regions followed by neuroblasts in the striatum (OBNBL3, 46.7%) and neuroblasts in the hippocampus (DGNBL1, 46.3%). The lowest proportion of cells represented in clones were observed for ependymal cells (EPEN, 1.3% to 4.9% of EPEN cells across all regions), TEGLU16 piriform pyramidal neurons (2.5%), and TEINH21 inhibitory neurons (7.8%) in the cortex. In line with our previous observation and as expected, we found a high correlation (r = 0.57) between the number of cells in clones and barcode expression level for each cell type (**Extended Data Fig. 14**). In conclusion, we captured clonal information about most cell types found across different regions of the mouse telencephalon and demonstrated that reconstruction of clonal relationships using TREX has a very low error rate.

### Regional distributions of clonally related cells

Because we isolated barcoded cells from cortex, striatum and hippocampus, we asked how often clonally related cells spread across these areas. By calculating the proportions of cells across each forebrain region for each cloneID, we observed that the cells of 1,880 clones (79.7%) accumulated in a single region (**Fig. 3a, b**). This corresponds to 648 clones found solely in cortex (CX, 27.5%), 656 clones in striatum (STR, 27.8%) and 576 clones in hippocampus (HC, 24.4%). Clonal dispersion of progenitors across more than one region was less frequent and was observed for 282 clones (11.9%) spreading across CX and STR as well as CX and HC (182 clones, 7.7%), but rarely between STR and HC (9 clones, 0.4%) or all three regions (7 clones, 0.3%). This indicates that most clonally related cells show limited regional dispersion across the mouse telencephalon.

To assess which cell types were associated with dispersed clones, we determined the cell type composition of clones spread across multiple forebrain regions relative to the total number of cells in clones for each cell type (**Fig. 3c, d, Extended Data Fig. 15**). We found that clonally related cells that crossed the CX/STR boundary often contained inhibitory neurons such as MGE-derived neurogliaform cells (TEINH16) in the CX (**Fig. 3e**). Inhibitory neurons shared a cloneID with medium spiny neurons (MSN1) and grey matter astrocytes (ACTE2) in the STR as well as neuronal intermediate progenitor cells (SZNBL) and oligodendrocyte subtypes in both STR and HC. These data suggest that MGE-derived cortical interneurons are generated by distinct progenitor cells and revealed that individual progenitors can give rise to both neurons and oligodendrocytes. Also, subventricular zone neural stem cells (RGSZ) in the STR often shared a cloneID with cells such as grey matter astrocytes (ACTE2), layer 2/3 excitatory neurons (TEGLU7) and all oligodendrocyte subtypes in the CX (**Fig. 3f**). This demonstrates a direct clonal relationship between E9.5 progenitor cells that generate RGSZ neural stem cells and those that produce neurons and glia cells for the other regions of the telencephalon during embryonic development. A similar observation has been made using viral DNA barcoding at E11.5 (ref^38^) and our results corroborate the authors predictions about an earlier clonal relationship between adult and embryonic neural stem cells.

Many cell types that were specifically found in the HC such as neural stem cells in the subgranular zone, neuroblasts, excitatory and granule neurons shared a cloneID with multiple other cell types in HC, but rarely with other cell types in CX indicating an early segregation of progenitor fields for both regions (**Extended Data Fig. 16a-d**). However, clones with Cajal-Retzius cells (CR) were an exception: these clones rarely contained other cell types and often shared a cloneID with CR cells in CX (**Fig. 3g**). We quantified the proportions of CR cells across both regions for each cloneID and observed that the 24.6% of cloneIDs accumulated in CX, 49.3% in HC and 26.1% spread across both CX and HC (**Extended Data Fig. 16e, f**). CR cells are among the first-born neurons critical for brain development and our data confirm that these cells originate from three distinct sites in the brain^39^ and further indicate that the progenitors from disparate embryonic fields converge in their differentiation to produce transcriptionally similar cells.

The anatomical boundary between HC and STR was very rarely crossed (**Fig. 3h**) and cell types associated with such clones were mostly glial cells of the oligodendrocyte lineage such as oligodendrocyte precursor cells (OPC) and committed oligodendrocyte precursors (COP1). This suggests that oligodendrocytes in both HC and STR are derived from a common progenitor most likely located in the ventral forebrain which generates OPCs that subsequently migrate widely into all parts of the telencephalon before differentiating^40^.

Finally, clonally related immune cells comprising microglia (MGL1, MGL3) and perivascular macrophages (PVM1) showed a widespread regional dispersion and crossed anatomical boundaries between CX, STR and HC 1.3-fold to 9-fold more often than neuroectoderm-derived clones (**Fig. 3i, j**). This suggests that myeloid progenitors and their progeny undergo extensive migration to populate large areas of the forebrain.

### Fate distributions of clonally related cells

We investigated the distribution of cloneIDs across cell types by calculating the proportions of cells within each major cell class (astroependymal, immune, projection neurons, interneurons, immature neurons, oligodendrocytes) for each cloneID. We found that immune cells (n = 84 clones) consisting of microglia and perivascular macrophages constitute a separate lineage as expected (**Fig. 4a**). Out of the remaining 2,276 neuroectoderm-derived clones, a total of 1,193 clones (52.4%) contained at least two different cell types, e.g. 430 clones (18.9%) contained both astroependymal cells and oligodendrocytes while 40 clones (1.8%) contained projection neurons, immature neurons, astroependymal cells as well as oligodendrocytes (**Fig. 4b**). The remaining 1,083 neuroectoderm-derived clones contained only one of the five major cell types and such clones were also observed among the largest clones (**Extended Data Fig. 17**). While this might suggest that lineage-restricted progenitor cells exist in the E9.5 mouse neuroepithelium, it is not possible to conclude that a strictly “uni-potential” progenitor was indeed present during barcoding, because only a small sample of its progeny had been isolated.

**Fig. 4.**
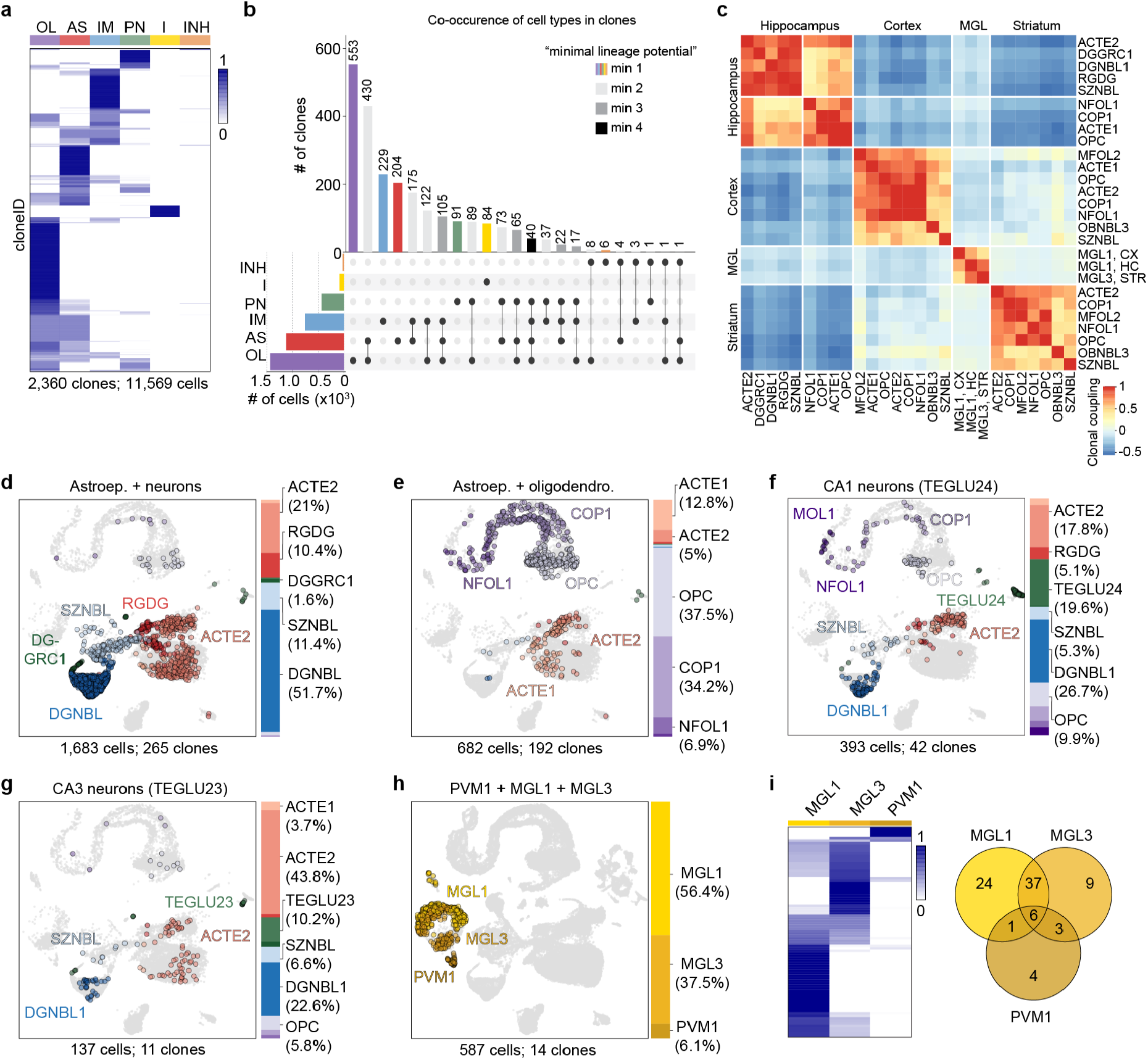
TREX reveals fate-biased progenitors of neuroectodermal and myeloid origin. **a**, Heatmap showing the proportions of each major cell type (columns) per cloneID (rows). For each cloneID, the proportions of cells within each cell type were calculated, scaled by row and coloured as shown in the figure. All 2,360 clones and a total of 11,569 cells were considered. OL, oligodendrocytes; AS, astroependymal cells; IM, immature neurons; PN, projection neurons; I, immune cells; INH, inhibitory interneurons. **b**, “UpSet” plot showing the co-occurrence of cell types in the same clone. All clones as shown in panel a were plotted. The bar plot shows the number of clones containing a particular combination of cell types and each bar is a different combination. The graphical table underneath the bar plot displays what those cell type combinations are. Each row is one cell type labelled as in panel a. For example, the most common subset of clones contains only OL cells (n = 553 clones, purple bar) followed by OL and AS cells (n = 430 clones, second light grey bar). The black dots and lines show the combination of cell types that make up each cluster or subset of clones. The smaller bar plot to the left of the graphical table shows the unconditional frequency count of each cell type across all subsets. We cannot exclude that a clone is indeed multipotential (= containing cells of more than one type) and the true number of multipotential clones is likely underestimated, because only a small sample of its progeny has been isolated. Therefore, we label clones as “minimally uni-potential” (colored bars), “minimally bi-potential” (light grey bars), “minimally tri-potential” (medium grey bars), “minimally tetra-potential” (dark grey bars). **c**, Heatmap showing correlation between clonal coupling scores defined as the number of shared cloneIDs relative to randomized data for each pair of cell types (see methods for details). High correlation values indicate a clonal relationship between cell types and a common progenitor cell. Clustered using complete linkage method. **d, e**, Precursor cells in the hippocampus neuroepithelium are biased to generate one of two fates. UMAP visualizations and bar plots for clones containing astroependymal (ACTE2, RGDG) and neuronal cells (DGNBL, SZNBL, DGGRC1) associated with fate 1 (d) as well as clones containing astroependymal cells (ACTE1, ACTE2) and oligodendrocyte subtypes (OPC, COP1, NFOL1) associated with fate 2 (e). **f, g**, Early fate specification of CA1 and CA3 excitatory neurons. Clones containing CA1 (TEGLU24) neurons never contained CA3 (TEGLU23) neurons (f) and vice versa (g). Otherwise these clones contained the same cell types including ACTE1, ACTE2, DGNBL1, SZNBL, OPC and COP1. **h, i**, Macrophages of the CNS parenchyma and CNS borders share the same precursor cell. UMAP visualization and bar plot for all 14 clones that contained perivascular macrophages (PVM1, h). Heatmap showing the proportion of cells classified as MGL1, MGL3 or PVM1 for each cloneID and Venn diagram displaying the number of clones containing cells of one, two or all three types (i) considering all 84 clones with 2,476 immune cells.

To systematically assess lineage relationships among subclasses of all cell types, we investigated the probability of recovering shared cloneIDs from all pairs of profiled cells in the mouse brain. We calculated the clonal coupling score defined as the number of shared cloneIDs relative to randomized data^23^ yielding values that range from positive (related cells) to negative (unrelated cells) for each brain (**Extended Data Fig. 18**). To summarize the data for all brains, we focused on the 27 cell types found in clones with at least 3 cells per clone across all four brains and determined the pairwise correlation between coupling scores. Hierarchical clustering of the pairwise correlations revealed four distinct groups of clonally related cells corresponding to diverse cell types of the cortex, hippocampus and striatum as well as microglia from all three regions (**Fig. 4c**). These results corroborated our previous observations regarding the limited clonal dispersion of most neuroectoderm-derived cell types across the mouse telencephalon.

We observed a strong clonal coupling in the hippocampus between neuronal and astroependymal cells (fate 1) as well as between astroependymal cells and oligodendrocytes (fate 2) indicating that these cells originate from two fate-biased pools of progenitor cells. We found that 265 clones containing 1,683 cells were biased towards fate 1 (**Fig. 4d**) and consisted mainly of neuronal cell types such as dentate gyrus neuroblasts (DGNBL1, 51.7%), neuronal intermediate progenitor cells (SZNBL, 11.4%) and granule neurons (DGGRC1, 1.6%) as well as astroependymal cells including grey matter (protoplasmic) astrocytes (ACTE2, 21%) and radial glia-like cells (RGDG, 10.4%). A total of 192 clones with 682 cells were biased towards fate 2 (**Fig. 4e**) and contained mainly oligodendrocyte subtypes such as oligodendrocyte precursor cells (OPC, 37.5%), committed oligodendrocyte precursors (COP1, 34.2%) as well as astroependymal cells including white matter (fibrous) astrocytes (ACTE1, 12.8%) and grey matter (protoplasmic) astrocytes (ACTE2, 5%). One population of progenitor cells, fate 1, likely corresponds to the embryonic precursors of adult neural stem cells^41^ that are biased to generate astroependymal cells and dentate granule neurons as early as E9.5. The second precursor cell population, fate 2, mainly contains oligodendrocyte subtypes and could represent a major source of hippocampal glia cells involved in myelin formation and maintenance.

We also investigated the cloneID distribution across cell types that were not included in the clonal coupling analysis, because they were not isolated from all four brains and/or they were not contained in clones with at least 3 cells per clone. Interestingly, we never observed hippocampal CA1 (TEGLU24) and CA3 (TEGLU23) excitatory neurons in the same clone that otherwise contained identical cell types (**Fig. 4f, g**). Because the number of clones containing at least one CA1 or CA3 neuron was small (42 clones with 77 CA1 cells and 11 clones with 14 CA3 cells), we cannot exclude that these cells share a common progenitor. However, our observations are in agreement with previous studies about the early specification of CA field identity^42^ and might indicate a fate specification (or at least fate bias) as early as E9.5.

We investigated the clonal relationships between microglia in the brain parenchyma (MGL1, MGL3) and perivascular macrophages (PVM1) located at CNS borders. We found that 10 out of 14 clones (n = 587 cells) that contained PVM1 cells also contained one or both microglia subtypes (**Fig. 4h, i**). Compared to 331 MGL1 cells (56.4%) and 220 MGL3 cells (37.5%), these clones contained only 36 PVM1 cells (6.1%). Because barcode expression levels and proportion of cells in clones were similar for MGL1, MGL3 and PVM1 (**Fig. 2g, Extended data Fig. 14**), this observation indicates that the common progenitor for all three cell types largely generates microglia and few perivascular macrophages. While it has been established that microglia are derived from mesodermal progenitors^43,44^, it has been shown only recently that the same early embryonic precursors also generate perivascular macrophages^45,46^. Our results are in line with this observation and further revealed that microglia are generated in much larger numbers than perivascular macrophages from a common progenitor cell.

### Spatial profiling of transcriptomes, cell types and clones

Next, we developed Space-TREX, a method based on spatial transcriptomics (ST)^26^ that enables simultaneous clonal tracing and expression profiling of barcoded mouse brain sections *in situ* (**Fig. 5a**). We introduced immunostaining of intracellular antigens into the protocol enabling combined profiling of spatial gene and protein expression together with clonal barcodes in the same tissue section (**Fig. 5b; Extended Data Fig. 19a-d**). Because ST relies on the capture of transcripts in spots with a diameter of 55 µm, most spots contain between 1-10 cells with an average of about 4 cells (**Extended Data Fig. 19e**). However, not every cell in the tissue is barcoded and out of all spots containing an EGFP+ cell, 81% of spots contain only one barcoded cell and the rest more than one barcoded cell (**Extended Data Fig. 19f**). Therefore, it can be assumed that a cloneID captured in a spot originates most often from a single barcoded cell and we can reveal its identity using protein expression data collected for the same section.

**Fig. 5.**
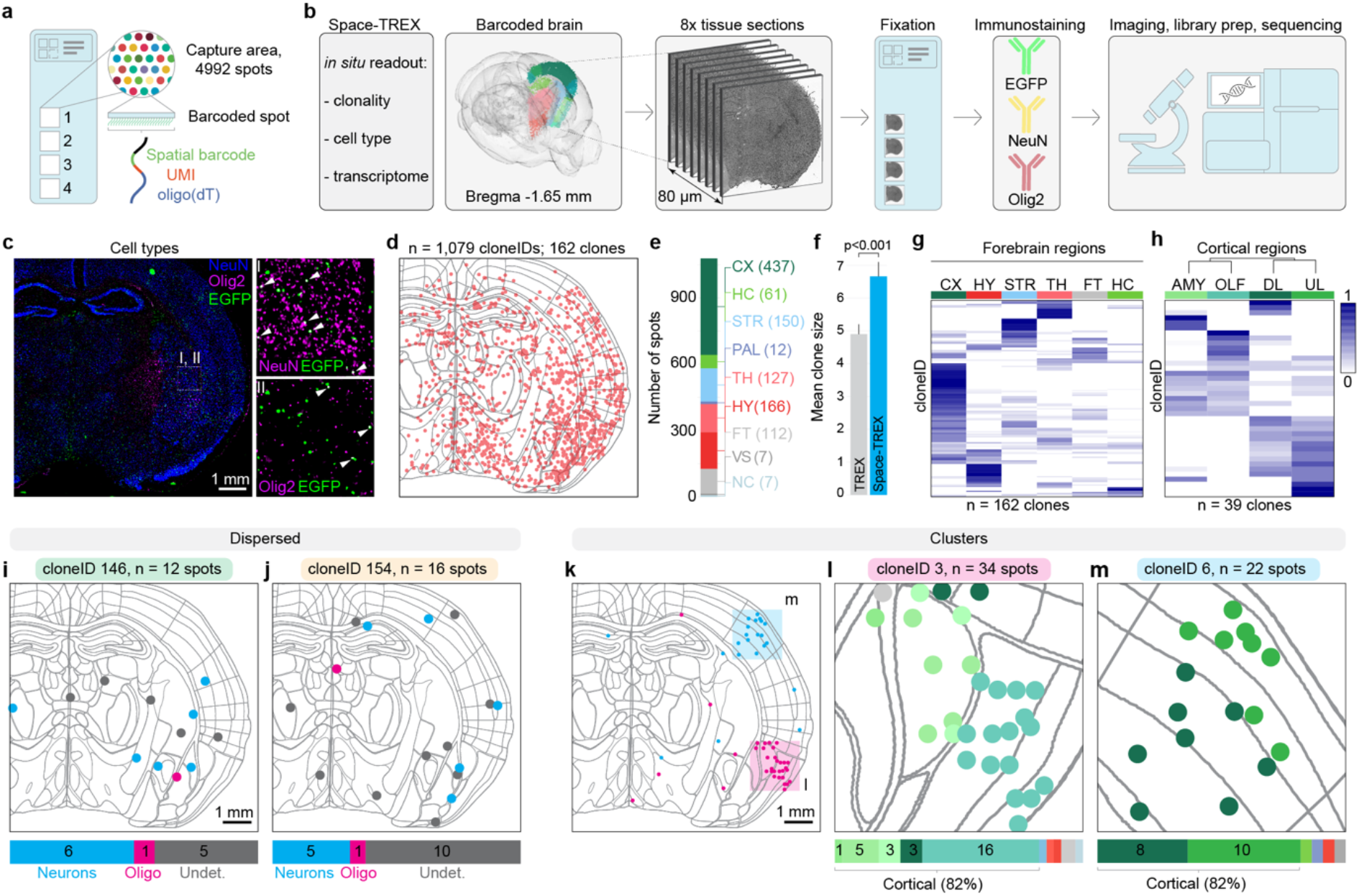
Space-TREX enables simultaneous profiling of transcriptomes and clones *in situ*. **a**, Glass slide layout used for spatial transcriptomics. Each slide consists of four 6.5 mm × 6.5 mm capture areas each containing 4,992 barcoded spots with a diameter of 55 µm and a center-to-center distance of 100 µm. Each spot contains spatially-barcoded capture oligonucleotides that bind mRNA released from the tissue enabling gene expression profiling *in situ*. **b**, Overview of the Space-TREX workflow. Adjacent 10 µm sections from a lentivirus-injected, barcoded brain were collected, fixed and incubated with fluorophore-conjugated antibodies recognizing EGFP (barcoded cells), NeuN (neurons) and Olig2 (oligodendrocytes) followed by imaging, library preparation and sequencing. **c**, Overview image of one immunostained brain section (left) and a zoom in showing barcoded neurons (I, white arrowheads) and barcoded oligodendrocytes (II, white arrowheads). **d**, A total of 1,321 cloneIDs were extracted from all analysed sections of which 1,079 cloneIDs were contained in 162 clones (defined as groups of two or more spots with the same cloneID). These cloneIDs are displayed as red dots projected onto a digital brain section containing anatomical reference outlines. **e**, Number of cloneID positive spots per major brain region. **f**, Clones reconstructed using Space-TREX (n = 162 clones) contain a significantly higher number of cells compared to TREX (n = 2,360 clones) which relies on tissue dissociation for scRNA-seq (Welch two sample t-test, t = 3.3205, df = 344.2, p-value = 0.0009948, 95% CI is 0.7 to 2.8). Values for clone sizes correspond to mean ± SEM. **g**, Regional distribution of all clonally related cells across forebrain region reveals patterns of clonal dispersion. Only regions containing more than 20 cloneIDs were considered resulting in the analysis of 162 clones with 1,053 spots. For each cloneID, the proportions of cells within each region were calculated, scaled by row and coloured as shown in the legend. CTX, cortex; HY, hypothalamus; STR, striatum; TH, thalamus; FT, fiber tracts; HC, hippocampus. **h**, High resolution spatial mapping of clonal dispersion across cortical regions reveals progenitors with a bias to generate cells of either amygdala (AMY) and olfactory areas (OLF) or upper layer (UL) and deep layer (DL) cells. Note that some cloneIDs were only found in UL or DL, respectively, suggesting fate bias of early progenitors. Clones containing at least 50% of clonally related cells in the displayed four regions were considered resulting in the analysis of 39 clones with 231 spots. **i, j**, Tangential migration of progenitors leads to widespread dispersion of neurons and oligodendrocytes. Two clones are shown as examples and clonally related spots are colour-coded by cell type (blue, neuron; magenta; oligodendrocyte; grey, undetermined). Note that regional information was left out for clarity and can be found the Extended Data Fig. 20a, b. **k-m** Radial migration of progenitors leads to clusters of clonally related cells illustrated with two clones (**k**) that mostly consist of cortical cells in the olfactory areas (**l**) and the primary somatosensory area (**m**). Clonally related spots are colour-coded by region with cortical regions in different shades of green. Note that cell type information as well as detailed regional annotation was left out for clarity and can be found in Extended Data Fig. 20c-e.

We hybridized 8 adjacent coronal sections from one P14 brain hemisphere barcoded at E9.5 covering the region about 1.65 mm posterior from Bregma and used antibodies targeted to EGFP, NeuN and Olig2 to identify barcoded cells, neurons and oligodendrocytes, respectively (**Fig. 5c**). To establish an integrated dataset containing information on spatial gene expression patterns, cell types, clones, and neuroanatomical definitions, we aligned brain sections to the Allen mouse brain reference atlas using an integrated computational framework^47,48^ (**Extended Data Fig. 19g-i**). The entire dataset after quality control contained information on the transcriptional profiles of 28,746 features that were distributed across all forebrain regions. We extracted a total of 1,321 cloneIDs of which 1,079 cloneIDs were contained in 162 clones (defined as groups of two or more spots containing the same cloneID) distributed across all sampled brain regions (**Fig. 5d, e; Extended Data Fig. 19j-n**). The number of cells per clone in the Space-TREX data (6.7 ± 0.4, mean ± SEM, n = 162 clones, **Fig. 5f**) was significantly larger than the clone size observed in the TREX data (4.9 ± 0.3, mean ± SEM, n = 2360 clones) indicating that cell loss leading to incomplete clones is reduced when using a spatial barcode readout (see also discussion).

Using the Space-TREX dataset, we investigated the regional dispersion of clonally related cells across the mouse telencephalon (**Fig. 5g**). In line with the TREX data, most clones showed a limited spread across all regions except for clones with cells located in white matter fiber tracts that are known be to be enriched for oligodendrocytes derived from highly migratory progenitors^40^. While most clonally related cells crossed boundaries of major anatomical regions at low frequencies, intra-regional dispersion was more common as observed for clones that mainly spread among cortical regions such as the amygdalar (AMY) and olfactory (OLF) areas as well as upper (UL) and deeper (DL) cortical layers (**Fig. 5h**). Interestingly, we observed extensive dispersion between either AMY/OLF or DL/UL suggesting that most early progenitor cells are restricted to generate cell types of either area but undergo more widespread migration within each area.

Next, we utilized cell type information available for barcoded cells and found that clones containing both neurons and oligodendrocytes show an extensive spread across multiple regions including cortical regions, striatum, pallidum, thalamus, hypothalamus and fiber tracts (**Fig. 5i, j; Extended Data Fig. 20a, b**). This mode of dispersion likely corresponds to tangential migration well described for interneurons^49^ that also share a common early progenitor with oligodendrocytes although these lineage relationships are not well understood^50^. We also observed neuronal clones that formed radially organized clusters mainly in the AMY/OLF areas as well as UL/DL areas of the cortex (**Fig. 5k-m; Extended Data Fig. 20c-e**). While a few members of these clones were more widespread, more than 80% of all clonally related cells were found in larger clusters spanning areas of around 1.75 mm x 1.75 mm. Interestingly, cells from dispersed clones were distributed across the dorsoventral axis within a single 10 µm section while cells from clustered clones were spread from the most anterior to the most posterior brain section spanning 80 µm (**Extended Data Fig. 20f-i**), Together, these data demonstrate that Space-TREX can be utilized for high-throughput mapping of clonal relationships and cell types *in situ* and we expect such an integrated approach to be highly relevant to relate gene expression, clonality as well as connectivity.

## Discussion

We have developed two complementary approaches that enable simultaneous clonal tracing and gene expression profiling of dissociated mouse brain cells using single-cell RNA-seq (TREX) and of entire mouse brain sections using spatial transcriptomics (Space-TREX). Using TREX, we unravelled that the regional dispersion of most clonally related cells is limited and that most dispersion occurs across the anatomical boundaries of cortex (CX) and striatum (STR) or cortex (CX) and hippocampus (HC), respectively. Often specific cell types are associated with dispersed clones such as interneuron subtypes (CX/STR), Cajal-Retzius cells (CX/HC) or oligodendrocyte subclasses (STR/HC). We discovered two fate-biased progenitor cell populations that exist as early as E9.5 in the hippocampal neuroepithelium and that tend to generate clones containing neuronal/astroependymal (fate 1) or astroependymal/oligodendrocyte (fate 2) cells, respectively. This finding suggests an unexpected early segregation of precursor cells and is consistent with fate 1 progenitors being the origin for Hopx+ precursors that continue to become adult neural stem cells in the mouse dentate gyrus^41^.

We unravelled unique features of myeloid-derived clones (microglia and perivascular macrophages) such as their large clone sizes and widespread dispersion across multiple forebrain regions compared to neuroectoderm-derived clones. The large clone size probably reflects the massive proliferation of brain macrophages required to colonize the entire CNS, because only a small number of precursors enters the brain before closure of the blood-brain barrier around E13 restricting access to immune cells that arise later in development^36^. It has been described that embryonic microglia migrate long distances within regions after entering the brain^51^ and we show that clonally related microglia also extensively migrate across anatomical boundaries to populate large areas of the brain. Both processes, microglia expansion and dispersion are central for brain homeostasis^52-54^, but remain only partially understood in particular at the clonal level. Thus, novel tools such as TREX enable systematic studies of the underlying molecular mechanisms within context of microglia clonality.

Using Space-TREX, we provide the first demonstration of high-density clonal tracking coupled to cell phenotyping and *in situ* sequencing of brain tissue and showed that patterns of clonal dispersion, tangential and radial migration, can be reconstructed using spatial transcriptomics.

Compared to previous approaches that utilize complex *in situ* hybridization schemes and fluorescence microscopy for barcode detection^14,17^, Space-TREX relies on widely available reagents and DNA sequencing thus enabling barcode readout in large tissue sections at scale^55^. Currently, (Space-)TREX is limited by undersampling due to loss of barcoded cells following isolation via FACS (35-64% of sorted cells recovered) and droplet encapsulation (50% of loaded cells recovered) as well as cloneID dropout from a subset of sequenced cells (24-51% contain a cloneID), resulting in clonal information for 2-7.5% of all initially isolated cells from a barcoded brain (**Extended Data Fig. 21**). Therefore, we expect that the true clone size is 13-to 50-fold higher than the average clone size observed under our experimental conditions and estimate that each neuroectoderm-derived clone contains between 52-200 cells while each myeloid-derived clone is composed of 390-1500 cells. While we, to our knowledge, report the first estimated clone sizes for myeloid-derived cells in the mouse brain, our clone size estimates for neuroectoderm-derived clones are in agreement with the recent finding that a cortical clone labelled at E9.5 contains 199 ± 29 cells (mean ± SEM, n = 13 clones)^56^.

The observed cell and barcode recovery rates are in line with other approaches employing a scRNA-seq readout of genetic barcodes in various model systems and highlight a general challenge for such methods (**Extended Data Fig. 22**). Thus, such approaches probably underestimate the true number of multipotent clones, but instead rely on sequencing thousands of cells to provide statistically robust insights about the fate bias of progenitor cells. One practical solution to reduce both cell and cloneID loss is using a plate-based assay with higher RNA detection sensitivity such as Smart-Seq3 (ref^57^). Further, undersampling could be completely eliminated by the adoption of a single-cell, high sensitivity *in situ* readout of cell types and barcodes using array-based spatial transcriptomics. Our Space-TREX data provide a first example of this: although the method currently lacks single-cell resolution and does not capture all cloneIDs, we found a significantly higher number of cells in clones compared to TREX.

Our *in vivo* barcoding approach is robust and can be applied to any tissue and cell type amenable to virus transduction without relying on a dedicated transgenic mouse line. Compared to classical fate mapping studies that rely on sparse labelling of cells in dozens to hundreds of animals, (Space-)TREX enables high throughput dense reconstruction of clonal relationships using only a few animals. In particular, our approach involves 13-31 times less animals than needed for typical fate mapping studies while recovering 3-13 times more clones in total and 98-182 times more clones on average per animal (**Extended Data Fig. 23**). In contrast to CRISPR-based lineage tracing systems^15,20,21,24,25^, our technology uses millions of diverse and compact barcodes that can be easily cloned as libraries enabling straightforward barcode readout and clone reconstruction. We used a constitutively expressed barcode, but also provide a Cre-inducible lentivirus for conditional barcode expression as well as a backbone containing an U6-gRNA expression cassette to couple clonal tracking with CRISPR-based perturbations (**Extended Data Fig. 24**). Other future modifications could include the insertion of cell type specific regulatory elements^58^, activity indicators^59^, optogenetic^60^ or chemogenetic^61^ probes for functional interrogation of clonally related cells^62,63^. Overall, we believe that an integrated approach such as Space-TREX is required to disentangle the complex relationships between cell identity, cell history and tissue anatomy required to understand both the healthy and diseased brain.

## Methods

### Plasmid cloning

LV-EF1a-H2B-EGFP (**Extended data Fig. 1a**) was based on LV-GFP^64^. To exchange the PGK1 promoter in LV-GFP with an EF1a promoter, the LV-H2B-EGFP backbone was amplified from LV-GFP using primer pair MRX1514/1515 (**Supplementary Table 1**) (IDT) and EF1a was amplified from plasmid dCas9-VP64_GFP^65^ using primer pair MRX1516/1517 (**Supplementary Table 1**). Both fragments were digested with SalI-HF and SpeI-HF (NEB) followed by T4 ligation (NEB) and transformation of NEB Stable cells (NEB). Plasmids were prepared from the transformants and verified via Sanger sequencing.

Additional reporter constructs (**Extended data Fig. 7j**) were cloned by exchanging EGFP from LV-EF1a-H2B-EGFP with TagBFP (Evrogen), TagRFP (Evrogen) or emiRFP670 (ref^66^). Fluorescent proteins were amplified from plasmids using the primer pairs listed in **Supplementary Table 1** (IDT), digested with BamHI/NsiI (NEB) followed by T4 ligation with digested backbone and transformation of NEB Stable cells. Plasmids were prepared from the transformants and verified via Sanger sequencing.

### Plasmid library construction and virus production

The backbone EF1-H2B-EGFP was digested with NsiI-HF and KpnI-HF (NEB) followed by heat inactivation, dephosphorylation using rSAP (NEB) and gel purification. An oligonucleotide library was prepared via low-cycle PCR using primer pair MRX1642/1643 (**Supplementary Table 1**) (IDT) on an ultramer MRX1680 (**Supplementary Table 1**) containing a random 30N sequence (cloneID) flanked by homology arms. The purified PCR product was inserted into digested backbone using Gibson assembly^67^ followed by ethanol precipitation and transformation of electrocompetent Endura cells (Lucigen). Bacteria were plated on a large 24.5 × 24.5 cm^2^ plate containing solid media and ampicillin and cells were scraped from the plate for plasmid DNA extraction using a plasmid midi kit (Qiagen) the next day. Lentiviral transfer plasmid libraries were used for virus particle production and titration by the core facility VirusTech at Karolinska Institutet using the packaging plasmids pMD2.G and psPAX2 (a gift from Didier Trono, Addgene plasmids # 12259 and # 12260) or sent to GEG-Tech (Paris, France). Viruses with titers of >10^9^ TU/ml were used for all applications.

### Next generation sequencing of lentivirus preparations

Viral RNA was isolated from 4 µl of a typical lentivirus preparation (∼2×10^6^ TU/µl) using the NucleoSpin RNA Virus mini kit (Macherey-Nagel). Purified RNA was used as a template for reverse transcription using the SuperScript VILO cDNA Synthesis Kit (Invitrogen) in a total volume of 80 µl following the manufacturer’s instructions (25°C for 10 min, 42°C for 60 min, 85°C for 5 min). The cDNA was equally divided into four separate PCRs for cloneID amplification and indexing using the primers in **Supplementary Table 1** and the PCR protocol in **Supplementary Table 2**.

The resulting libraries were sequenced on an Illumina NextSeq (**Supplementary Table 3**), aligned against a reference containing the 30 bp cloneID and flanking regions using the BWA-MEM algorithm^68^. A custom BASH script was used to extract unique cloneIDs and corresponding read counts.

### Estimating the fraction of uniquely labelled cells

First, we calculated the total number of cells at the timepoint of injection. If *N*_*t*1_ is the number of labelled cells at E11.5, Δ*t* is the time difference in days and *f* is the frequency of cell divisions per day, then the number of transduced cells *N*_*t*0_ is:

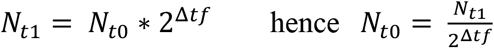

We determined *N*_*t*1_ = 41,450 cells, Δ*t* = 2 days and *f* = 2 divisions/day (ref^30^) thus *N*_*t*1_ = 2,591 ± 220 cells, or approximately 2,600 as noted in the main text. Second, we estimated the fraction of uniquely labelled cells. For a number of uniformly distributed barcodes (*N*) and a small number of used barcodes (*k*) to label progenitor cells, the fraction F of uniquely labelled cells can be approximated as:

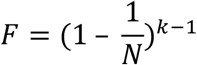

However, the observed distribution of barcode abundance is not perfectly uniform in our library, which means that cells are more likely to be labelled with some barcodes than with others. As discussed extensively in previous studies^55,69^, the expected number of non-uniquely labelled cells, *E*(*X*), is then given by:

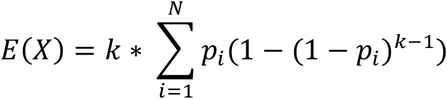

where *p*_*i*_ is the probability of picking the cloneID *i* = 1… *N, k* is the number of infected progenitor cells and *N* is the total number of injected cloneIDs. We typically injected *N* = 0.94 × 10^6^ cloneIDs and estimated *k* = 2,591 cells, implying *E*(*X*) ≈ 11 non-uniquely labelled cells. This corresponds to 0.41% of non-uniquely labelled cells or about 99.6% uniquely labelled cells as stated in the main text.

### Mice

CD-1 mice obtained from Charles River Germany were used for all experiments. Animals were housed in standard housing conditions with 12:12-hour light:dark cycles with food and water ad libitum. All experimental procedures were approved by the Stockholms Norra Djurförsöksetiska Nämnd.

### Ultrasound-guided *in utero* microinjection

To target the developing mouse nervous system, a modified version of a previously published procedure^64^ was used. Briefly, timed pregnancies were set up overnight, plug positive females were identified the next morning and counted as embryonic (E) age 0.5. Ultrasound check was performed at E8.5 to verify the pregnancy. Pregnant females at E9.5 of gestation were anaesthetized with isoflurane, uterine horns were exposed, each embryonic forebrain injected with 0.6 µl of lentivirus and 4-8 embryos injected per litter. Surgical procedures were limited to 30 min to maximize survival rates.

### Immunostaining and imaging of embryonic and postnatal tissue

Embryonic (E) age 11.5 mouse embryos were collected in ice-cold PBS, fixed in fresh 4% formaldehyde (FA) overnight at 4°C, placed in 30% sucrose overnight at 4°C and embedded in Tissue-Tek® O.C.T.™ (Sakura). 20 µm thick sections were cut using a NX70 Cryostat (ThermoFisher Scientific) and thaw-mounted onto Superfrost Plus™ microscope slides (ThermoFisher Scientific). Postnatal mice were sacrificed by isoflurane overdose followed by transcardial perfusion with ice cold PBS followed by 4% FA. Brains were postfixed in 4% FA overnight and 50 µm sections prepared using a VT1000S vibratome (Leica). Frozen and free-floating tissue sections were incubated with blocking/permeabilization buffer (5% donkey serum and 0.3% Triton X-100 in DPBS) and immunostained with antibodies against EGFP (chicken, 1:2000, Aves Labs, AB_2307313), NeuN (rabbit, 1:500, Atlas Antibodies, AB_10602305), Sox9 (goat, 1:300, R&D Systems, AB_2194160), Sox10 (goat, 1:300, R&D Systems, AB_442208) or Iba1 (rabbit, 1:500, Wako, AB_839504) at 4°C overnight. Sections were washed three times with DPBS and incubated with fluorophore-conjugated secondary antibodies (1:500) and DAPI (1 µg/ml) in blocking buffer at room temperature for one hour. Sections were washed three times with DPBS and mounted in ProLong® Diamond Antifade Mountant (ThermoFisher Scientific). Confocal images were captured using a laser scanning confocal microscope (LSM700, Carl Zeiss) using a Plan-Apochromat 10x/0.45 or 20x/0.8 objective. Image processing and analysis was performed using the software Fiji^70^.

### Single-cell dissociations and flow cytometry

Mice were sacrificed with an overdose of isoflurane, followed by transcardial perfusion with ice cold artificial cerebrospinal fluid (aCSF: 87 mM NaCl, 2.5 mM KCl, 1.25 mM NaH_2_ PO_4_, 26 mM NaHCO_3_, 75 mM sucrose, 20 mM glucose, 2 mM CaCl_2_, 2 mM MgSO_4_). Mice were decapitated, the brain was collected in ice-cold aCSF, 1 mm coronal slices collected using an acrylic brain matrix for mouse (World Precision Instruments) and the regions of interest microdissected under a stereo microscope with a cooled platform. Tissue pieces were dissociated using the Papain dissociation system (Worthington Biochemical) with an enzymatic digestion step of 20-30 min followed by manual trituration using fire polished Pasteur pipettes. Dissociated tissue pieces were filtered through a sterile 30 µm aCSF-equilibrated Filcon strainer (BD Biosciences) into a 15 ml centrifuge tube containing 9 ml of aCSF and 0.5% BSA. The suspension was mixed well, cells were pelleted in a cooled centrifuge at 300g for 5 min, supernatant carefully removed, and cells resuspended in 1 ml aCSF containing reconstituted ovomucoid protease inhibitor with bovine serum albumin. A discontinuous density gradient was prepared by carefully overlaying 2 ml undiluted albumin-inhibitor solution with 1 ml of cell suspension followed by centrifugation at 100g for 6 minutes at 4°C. The supernatant was carefully removed, the cell pellet resuspended in 1 ml aCSF containing 0.5% BSA and the cell suspension transferred to a round bottom tube (BD Biosciences) for flow cytometry. Single EGFP+ cells were sorted on a BD Influx equipped with a 140 µm nozzle and a cooling unit with a sample temperature of 4°C and collected into a DNA LoBind tube (Eppendorf) containing aCSF with 0.5% BSA. All EGFP+ cells per sample were sorted and pelleted in a cooled centrifuge at 300g for 5 min. The supernatant was carefully removed, the cell pellet resuspended in a minimal volume of aCSF and the cell concentration determined using a Bürker chamber. Importantly, aCSF equilibrated in 95% O2/ 5% CO2 was used in all steps, and cells were kept on ice or at 4 °C at all times except for enzymatic digestion.

### Bulk profiling of barcoded brain cells

To assess the number of successfully integrated barcodes in a typical experiment we isolated genomic DNA from transduced cells for bulk sequencing of the cloneID locus. Briefly, E9.5 mouse embryonic forebrains were injected as described above, embryos were collected at E15.5, decapitated and brains transferred into cold 1x HBSS on ice. Whole embryonic brains were cut into small pieces and processed as described above with the exception of a shorter enzymatic digestion step of 10 min. About 1,000 EGFP+ cells were sorted on a BD Influx (see above), collected into a 96-well plate containing 5.6 µl cell lysis buffer, followed by a 10-minute incubation step on ice and addition of 2.8 µl neutralization buffer. The entire lysate was used for PCR amplification and indexing of cloneIDs using the primers in **Supplementary Table 1** and the PCR protocol in **Supplementary Table 2**.

The resulting libraries were sequenced on an Illumina NextSeq (**Supplementary Table 3**), sequencing reads were aligned against a reference containing the 30 bp cloneID and flanking regions using the BWA-MEM algorithm^68^. A custom BASH script was used to extract unique cloneIDs with corresponding read counts and cloneIDs with reads in 0.9 quantile were kept.

### Single-cell RNA-seq

Two brains (brains 1-2) were processed using the 10X Genomics Chromium Single Cell Kit Version 2 (v2) and three brains (brains 3-5) using the 10X Genomics Chromium Single Cell Kit Version 3 (v3) (**Supplementary Table 3**). Suspensions from barcoded brains were prepared as described above, counted and resuspended aCSF and added to 10x Chromium RT mix. Suspensions from control brains prepared as described above and diluted in aCSF to concentrations between 800-1000 cells/µl and added to 10x Chromium RT mix. For downstream cDNA synthesis (12 PCR cycles), library preparation, and sequencing, we followed the manufacturer’s instructions.

### Data normalization and cell filtering for single-cell RNA-seq

Overall, three regions from four barcoded brains and from one control brain were sequenced using 10X Chromium v2 or v3. Because the number of cells per region for the control brain were much higher than the corresponding number of cells for any barcoded brain (**Supplementary Table 3**), we downsampled the control datasets to about 9,000 cells (cortex), 8,000 cells (hippocampus) and 7,000 cells (striatum). The gene expression matrices obtained after running cellranger count were merged by region (cortex, striatum, hippocampus) using merge() in Seurat v3 (ref^71^). All genes expressed in ∼0.1% of all cells were kept and all cells expressing 500 to 10000 genes were kept in the merged data. The data were log-normalized with a scale factor of 10000 using the NormalizeData() function followed by linear transformation (scaling) of data. Doublet removal was done using mutually exclusive markers for various cell types (Igf2, Pf4, Hexb, Rsph1, Pdgfra, Bmp4, Mog, Clic6, Rgs5, Cldn5, Reln, Igfbpl1, Slc32a1, Slc17a7, Aldoc). A cell cycle score was assigned to each cell and the difference between the G2M and S phase scores was regressed out. Highly variable features were selected using FindVariableFeatures() followed by PCA and the use of significant PCs (usually 10-30) for graph-based clustering (SNN graph calculation and clustering using Louvain). After determining differentially expressed genes, we manually assigned major cell classes to each cluster (Astroependymal, Immune, Neurons, Oligodendrocytes, Vascular) using canonical markers. We then split cells by major cell type, performed subclustering and extensively annotated each cluster based on canonical marker genes from published data and from www.mousebrain.org^3^. At each step, we removed (1) clusters classified with ambiguous labels and (2) outlier cells on the fringes of clusters in UMAP space to further eliminate doublets. We annotated clusters using the same mnemonic identifiers as provided on www.mousebrain.org and added corresponding cell type location and general description as metadata. Finally, we merged all cells into a single file together with metadata and annotations. The filtered cellIDs were exported and used as input for cloneID extraction and clone calling using the TREX Python pipeline (see below). Following clone calling, the obtained cloneIDs were added as metadata to each Seurat object.

### Biological pathway analysis between barcoded and control samples

Lentivirus vectors are derived from the human immunodeficiency virus type 1 (HIV-1)^72^. To investigate the effect of lentivirus transduction on cellular physiology, we analysed genes expressed during virus infection which included 195 genes involved in TLR signalling, TNF signalling, chemokine signalling, MHC presentation, cell cycle and apoptosis (KEGG pathway: mmu05170). Because our dataset contained an imbalanced number of cells per major cell type (astroependymal, immune, neurons, oligodendrocytes) and condition (non-injected control, lentivirus/barcoded), we downsampled the dataset such that each cell type per condition (e.g. “neurons_control” and “neurons_lenti”) contained an equal number of cells. We plotted expression values of non-zero expressed genes (n = 195) related to virus infection for single cells as heatmaps grouped by condition or major cell type. For each cell type, we analysed differentially expressed between both conditions (logfc.threshold ≥ 1) on normalized and variance stabilized downsampled datasets.

### CloneID enrichment from cDNA for single-cell RNA-seq

A nested PCR strategy on full length cDNA obtained during the first steps of 10X Genomics Chromium Single Cell Kit Version 3 library preparation was employed for enrichment of cloneIDs (**Extended data Fig. 7a**) using the primers listed in **Supplementary Table 1**. Briefly, full length cDNA was used as a template for PCR1 with primer pair MRX1587/14 followed by purification and PCR2 on the purified product with primer pair MRX1587/1588. Finally, 1% of the purified PCR2 product was used for indexing with MRX1589 and a unique i7 index primer from the Chromium i7 Multiplex Kit (10X Genomics) (**Supplementary Table 1-3**). Each amplicon library was sequenced on a MiSeq or NovaSeq6000 (**Supplementary Table 3**). We used Cell Ranger count for data processing of amplicon libraries and the TREX (see below) to extract cloneIDs as done for transcriptome expression libraries.

### Extraction of cloneIDs and clone calling for single-cell RNA-seq

Raw 10X Genomics Chromium Version 2 or 3 sequencing data were preprocessed with Cell Ranger v3.0.1. As reference for read mapping, Cell Ranger was configured to use a custom reference consisting of the GRCm38 (mm10) genome and an additional sequence representing the H2B-EGFP-N transgene, in which the cloneID region was marked with “N” wildcard characters. The resulting BAM file of aligned sequencing reads was then processed with TREX, our custom Python tool for cloneID extraction and clone calling. TREX uses only reads from filtered cells (see above) that align to the H2B-EGFP-N transgene. CloneIDs are recovered from those alignments that cover the masked cloneID region. If soft clipping is encountered at one of the bases adjacent to the region, the alignment is assumed to continue ungapped into the region. All cloneIDs with identical UMIs that come from the same cell (have the same cellID) are collapsed to a consensus sequence. To error-correct cloneIDs, they are single-linkage clustered using a Hamming distance of at most five as linking criterion. In each cluster, all its cloneIDs are replaced with the cloneID occurring most frequently in that cluster. From the resulting final cellID-cloneID combinations, those that are only supported by one UMI and one read are discarded. Also removed are cloneIDs that are supported by only one UMI and have a high frequency in another cell. We assume that those cloneIDs are contaminations.

The cleaned data are transformed into a count matrix showing UMI counts for each cloneID in each cell. This matrix is used to sort cells into clones of cells with similar cloneID combinations. Clonally related cells were identified as described previously^18^. Briefly, the Jaccard similarity between each pair of cloneID expressing cells was calculated using the R package proxy^73^. A Jaccard score of 0.7 was used as a cut-off for related cells and clones were defined as groups of two or more related cells. CloneIDs were added as metadata to each dataset.

### Calculation of clonal coupling scores

For each brain we calculated clonal coupling scores defined as the number of shared cloneIDs relative to randomized data^23^ considering all clones containing at least 3 cells per clone. We randomized the clone-cell type associations, while preserving the number of cell types related to each clone, and the number of clones related to each cell type, to create 1000 randomized data sets^74^. This allowed us to estimate the number of overlaps expected to occur by chance. We compared the observed clonal data to randomized data sets to obtain empirical p-values and z-scores. These indicate, for each pair of cell types, how often we expect to see the observed clonal association yielding values that range from positive (clonally related cells) to negative (clonally unrelated cells). To summarize the clonal coupling scores for four brains, we kept only cell types found in clones in all brains. For each brain, the Pearson correlations of z-scores between each pair of cell types were calculated, the correlation coefficients were transformed using Fisher Z-transformation and averaged to represent clonal coupling scores for all brains.

### Tissue processing and library preparation for Spatial Transcriptomics

Mice were sacrificed with an overdose of isoflurane, followed by transcardial perfusion with ice cold artificial cerebrospinal fluid (aCSF: 87 mM NaCl, 2.5 mM KCl, 1.25 mM NaH_2_ PO_4_, 26 mM NaHCO_3_, 75 mM sucrose, 20 mM glucose, 2 mM CaCl_2_, 2 mM MgSO_4_). Mice were decapitated, the brain was collected in ice-cold aCSF, transferred to ice-cold Tissue-Tek® O.C.T.™ (Sakura) and snap frozen at -40°C in a bath of isopentane and dry ice. Eight consecutive 10 µm sections around AP -1.65 mm from Bregma were collected for processing using the 10X Genomics Visium Spatial Gene Expression kit.

The first four sections (V9-V12) were fixed in ice-cold methanol followed by rapid imaging (< 15 min for all sections) of EGFP and transmitted light signal using an epifluorescence microscope (Axio Imager.Z2, Carl Zeiss) equipped with a Plan-Neofluar 10x/0.3 M27 objective before further processing following the manufacturer’s instructions. The remaining four sections (V13-V16) were fixed in ice-cold methanol, briefly rinsed with DPBS, incubated with DPBS containing DAPI (1 µg/ml), FluoTag®-X4 anti-GFP conjugated to Atto488 (1:200, NanoTag Biotechnologies), NeuN-Alexa568 (rabbit, 1:400, Abcam, ab207282), Olig2-Alexa647 (rabbit, 1:200, Abcam, ab225100) and RNaseOUT™ (1 U/µl) at room temperature for 10 min. The sections were washed two times for 1 min with DPBS containing RNaseOUT™ (1 U/µl), mounted in 85% glycerol containing RNaseOUT™ (1 U/µl) and images were captured for all four fluorescent channels as well as the transmitted light channel. The coverslip was removed by immersing the slide in water, the slide was dried for 5 min at 37°C and further processed following the manufacturer’s instructions starting with the tissue permeabilization step.

### Data and image analysis for Spatial Transcriptomics

For each coronal brain section, the registered microscope image was used for manual alignment and tissue detection using the Visium Manual Alignment Wizard (10X Genomics) followed by running Space Ranger to obtain a gene expression matrix for each section (**Supplementary Table 3**). Each dataset was separately processed in Seurat v3 (ref.^71^) and only spots that expressed at least 300 genes were kept. We used SCTransform^75^ to normalize the data and detect high-variance features followed by dimensionality reduction, clustering, visualization in UMAP space and identification of spatially variable gene expression. The transcriptomic profiles of capture spots in each dataset were integrated with the merged v3 single-cell RNA-seq dataset to predict the underlying composition of cell types. Finally, all datasets were merged, spot IDs were exported and used as input for cloneID extraction and clone calling using TREX.

Fluorescent images acquired for four sections (V13-V16) were processed in R using a custom segmentation workflow (https://github.com/ludvigla/TREXSeg) that entails (1) 2D FFT convolution filtering, (2) image correction, (3) thresholding, (4) removal of speckles or other abnormal shapes and (5) watershedding to identify and label cells. The segmentation workflow was applied to each of three channels EGFP (barcoded cells), NeuN (neurons) and Olig2 (oligodendrocytes). To find co-localizing signals across two channels A and B, an overlap score was estimated for all pairs of nuclei (*i, j*) as intersect(A_i_, B_j_)/min(A_i_, B_j_) where A_i_ and B_j_ are the sets of pixels defining nuclei *i* and *j*. An overlap score of at least 50% was used to determine if the signal originated from the same nuclei. For alignment of all four sections, we used a manual image registration method implemented in the ManualAlignImages function from the STUtility package^76^. Briefly, the raw NeuN images were first masked distinguish the tissue from the background and to detect the tissue edges. The detected tissue edges were then manually rotated or shifted using the manual registration tool to fit the image and spot coordinates images V14-V16 to the reference image V13. All capture spot coordinates from V14-V16 were transformed to align with the coordinate system of V13 using the learned transformation functions. The same transformations were applied to the coordinates of the previously segmented nuclei in V14-V16. We calculated the pairwise 2D Euclidean distances between aligned spots and nuclei and selected the cell with shortest distance to the centroid position of each cloneID+ spot for assignment of cell type identity. For alignment of all four H&E stained sections (V9-V12), we used an automated image registration method implemented in the AlignImages function from STUtility with image V9 as reference. Similar to the manual registration method described above, the automatic registration uses masked H&E images to find a transformation function that minimizes the difference between the pixels outlining the tissue edges which is solved using an Iterative Closest Point (ICP) algorithm. All capture spot coordinates from V10-V12 were transformed to align with the coordinate system of V9 using the learned transformation function.

Registration of aligned images of brain tissue sections to the standardized Allen Mouse Brain Atlas (ABA) was done using WholeBrain^48^. We used an extended and inverted version of the H&E target image V9 and the NeuN target image V13 with Bregma coordinates AP -1.65 mm for registration of an entire brain section to the ABA. Anatomical information and ABA color codes were added to the merged Spatial Transcriptomics dataset and plotted using custom functions.

### CloneID enrichment from cDNA for Spatial Transcriptomics

A nested PCR strategy on full length cDNA obtained during the first steps of 10X Genomics Visium Spatial Gene Expression library preparation was employed for enrichment of cloneIDs using the primers listed in **Supplementary Table 1**. Briefly, unfragmented cDNA was used as a template for PCR1 with primer pair MRX1587/14 followed by purification and PCR2 on the purified product with primer pair MRX1587/1805 (**Supplementary Table 1-2**). Finally, 1% of the purified PCR2 product was used for dual indexing with Visium Dual Index primers (10X Genomics) (**Supplementary Table 3**). Each amplicon library was sequenced on a NextSeq (**Supplementary Table 3**).

### Extraction of cloneIDs and clone calling for Spatial Transcriptomics

Raw 10X Genomics Visium sequencing data from transcriptome and amplicon libraries (**Supplementary Table 3**) were preprocessed with Space Ranger v1.0. The pipeline included an alignment step where sequencing reads were mapped to the mm10 transcriptome and to an additional chromosome representing the H2B-EGFP-N transgene. Aligned sequencing reads of all spots were subjected to TREX for cloneID extraction as described above. From the resulting final spotID-cloneID combinations, only those cloneIDs supported by at least one UMI and more than one read are kept. Clone calling was performed based on Jaccard similarity as described above and cloneIDs were added as metadata to each dataset in Seurat.

### Overview of software and packages used in this study

The following and R software^77^ packages and libraries were used: BiRewire^74^, cowplot^78^, dplyr^79^, EBimage^80^, eulerr^81^, ggplot2 (ref.^82^), magick^83^, magrittr^84^, Matrix^85^, pheatmap^86^, proxy^73^, RColorBrewer^87^, reshape2 (ref.^88^), rgl^89^, SDMTools^90^, Seurat^75^, STutility^76^, tidyverse^91^, umap^92^, wholebrain^48^, zeallot^93^. The following Python software (https://www.python.org) packages and libraries were used: AmpUMI^94^, Loompy (http://loompy.org), NumPy^95^, Pandas^96^, Pysam (https://github.com/pysam-developers/pysam), Tinyalign (https://github.com/marcelm/tinyalign), xopen (https://github.com/marcelm/xopen).

## Data availability

All RNA sequencing datasets generated in this study are deposited in the Gene Expression Omnibus (GEO) under accession code GSE153424. All processed single-cell and spatial transcriptomics datasets are available as RDS files using link https://kise-my.sharepoint.com/:f:/g/personal/michael_ratz_ki_se/EndBZ9VI_rRHmHzZxrAwSZQBeE9e4RNmktbuCcHir1a5qQ?e=Ge2Fqm and password 8RMG.xbzH?3v9Ef4.

## Code availability

TREX source code is available under the MIT license from https://github.com/frisen-lab/TREX. Image segmentation and alignment code is available under the MIT license from https://github.com/ludvigla/TREXSeg.

## Author contributions

MR led experimental work, generated and analysed data and prepared figures. LvB developed the initial barcode extraction pipeline with help from GLM, analysed data, and helped with experiments supervised by MR. LL developed the cell segmentation and image alignment pipeline. MM and LvB further improved and expanded functionality of the barcode extraction pipeline. JOW developed the pipeline for assessing clonal coupling. JF and JL supervised the study. MR and JF wrote the manuscript with input from all authors.

## Acknowledgments

We thank Enric Llorens-Bobadilla, Igor Adameyko, Robert Harris, Konstantinos Meletis and Daniel Sheward for critical discussions; Lola Buades for help with drawing Figure 1a and Extended Data Fig. 2e; Sarantis Giatrellis for FACS assistance; Maria Genander and Christina Kantzer for the LV-GFP construct; Ilaria Testa for the emiRFP670 plasmid; the lab of Rickard Sandberg for access to NextSeq; the National Genomics Infrastructure in Stockholm funded by Science for Life Laboratory, the Knut and Alice Wallenberg Foundation and the Swedish Research Council for sequencing services; SNIC/Uppsala Multidisciplinary Centre for Advanced Computational Science for assistance with massively parallel sequencing and access to the UPPMAX computational infrastructure; ultrasound-guided *in utero* injections were performed by the Infinigene core facility and lentivirus production was done by the VirusTech core facility at Karolinska Institutet or GEG-Tech (Paris, France). JOW and MM are financially supported by the Knut and Alice Wallenberg Foundation as part of the National Bioinformatics Infrastructure Sweden at SciLifeLab. This study was supported by grants from the Swedish Research Council, the Swedish Cancer Society, the Swedish Foundation for Strategic Research, Knut och Alice Wallenbergs Stiftelse and the ERC to JF and a DFG Research Fellowship (RA 2889/1-1) to MR.

