## Extended Data for "Cell types and clonal relations in the mouse brain revealed by single-cell and spatial transcriptomics"

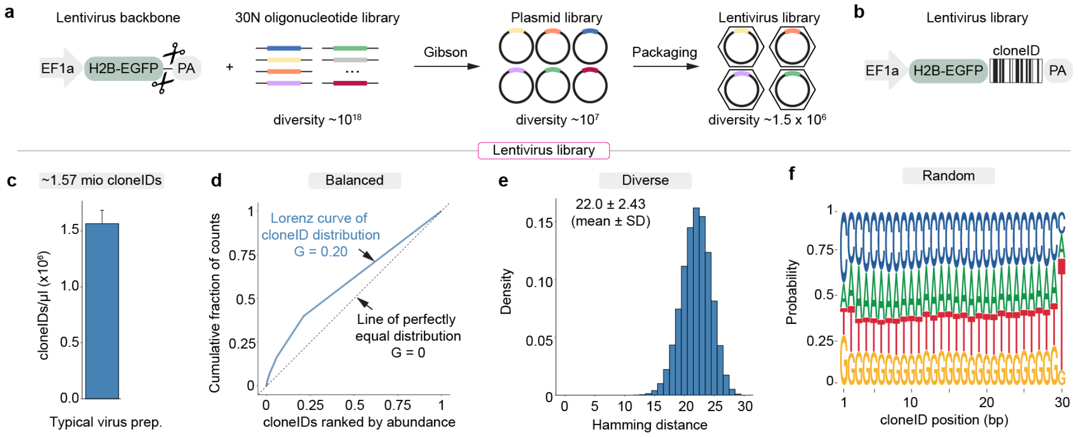


**Extended Data Fig. 1 | Generation and characterization of cloneID barcode libraries.** **a**, A lentiviral backbone containing a strong human elongation factor 1 alpha (EF1a) promoter driving the expression of nuclear-localized enhanced green fluorescent protein via fusion to histone 2b (H2B-EGFP) was constructed based on LV-GFP[1]. The EF1a-H2B-EGFP backbone was used for insertion of an amplified random 30N oligonucleotide library. The resulting plasmid library was subjected to lentiviral packaging to generate a lentivirus library with high functional titers greater than $10^9$ TU/ml. **b**, Each plasmid of the final library contains a unique random 30N barcode or cloneID downstream of the EGFP stop codon and known flanking sequences. **c-f**, Next generation sequencing was used for characterization of cloneID sequences in a typical lentivirus preparation (**g-f**). **c,** A total of 1,566,516 ± 120,080 cloneIDs/µl (mean ± SD, n = 4 virus preparations) were found in a typical lentivirus preparation. **d,** Lorenz curves (blue) were used to describe the inequality in the distribution of counts for all cloneIDs compared to the line of a perfectly equal distribution (dashed). The Gini coefficient (G), an overall measure of cloneID count inequality, is low for a typical lentivirus preparation indicating a uniform cloneID representation. **e,** The pairwise Hamming distance between all sequences for a random sample of cloneIDs (n = 10,000) was calculated for a representative lentivirus preparation demonstrating high cloneID sequence diversity. **f,** Sequence logo plot for a random sample of cloneIDs (n = 100,000) demonstrating that each cloneID is a random sequence based on similar probabilities for the occurrence of each of the four nucleotides (A, T, C, G) in the cloneID sequence at each of the 30 positions.

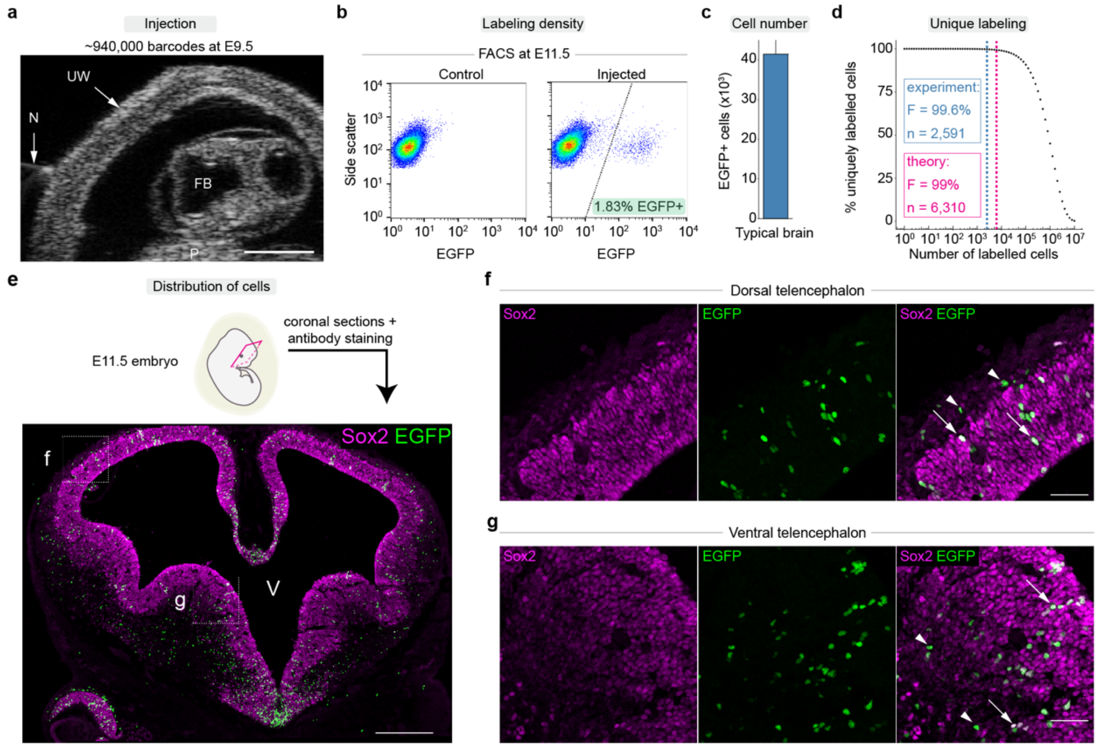


**Extended Data Fig. 2 | Unique labeling of mouse brain progenitors *in vivo*.** **a**, Ultrasound image of mouse embryo (dorsal view) at embryonic day (E) 9.5 showing injection needle (N), uterine wall (UW), placenta (P) and forebrain (FB, injection target site). We typically injected a lentivirus volume of 0.6 µl corresponding to about 940,000 unique cloneIDs. Scale bar, 1 mm. **b**, Fluorescent activated cell sorting (FACS) of single cell suspensions prepared from a non-transduced control (left) and a representative lentivirus transduced (right) mouse brain two days after injection at E11.5. The fraction of EGFP+ cells upon lentivirus-mediated barcoding was 1.83 ± 0.25% (mean ± SD, n = 3 brains). **c**, Barplot showing the total number of barcoded EGFP+ cells two days after injection at E11.5. We counted a total of 2,279,750 ± 259,650 (mean ± SD, n = 3 brains) cells in single cell suspensions prepared from a E11.5 mouse brain. Based on the fraction of labelled cells it can be estimated that a total of 41,450 ± 3,513 (mean ± SD, n = 3 brains) barcoded EGFP+ cells are present in a E11.5 mouse brain after virus injection at E9.5. **d**, Technically, the number of cloneIDs is sufficient to label 6,310 cells in a typical virus injection experiment (V = 0.6 µl, 0.94 x $10^6$ cloneIDs) with <1% barcode overlap between clones (magenta dashed line). Practically, around 2,600 cells are labelled in a typical experiment (blue dashed line) hence 99.6% of these cells were uniquely labelled with a cloneID. **e**, Top, Schematic of E11.5 mouse embryo used for cryosectioning following lentivirus injection into the E9.5 brain. Bottom, Representative image of a coronal E11.5 brain section showing Sox2+ neuronal progenitor cells located primarily in the ventricular zone as well as lentivirus-transduced EGFP+ cells distributed throughout the neural tube tissue. Scale bar, 500 µm. V, ventricle. **f, g,** Sox2+/EGFP+ double positive progenitor cells (arrows) and their Sox2-/EGFP+ daughter cells (arrow heads) located in the dorsal (**f**) and ventral (**g**) telencephalon, respectively.

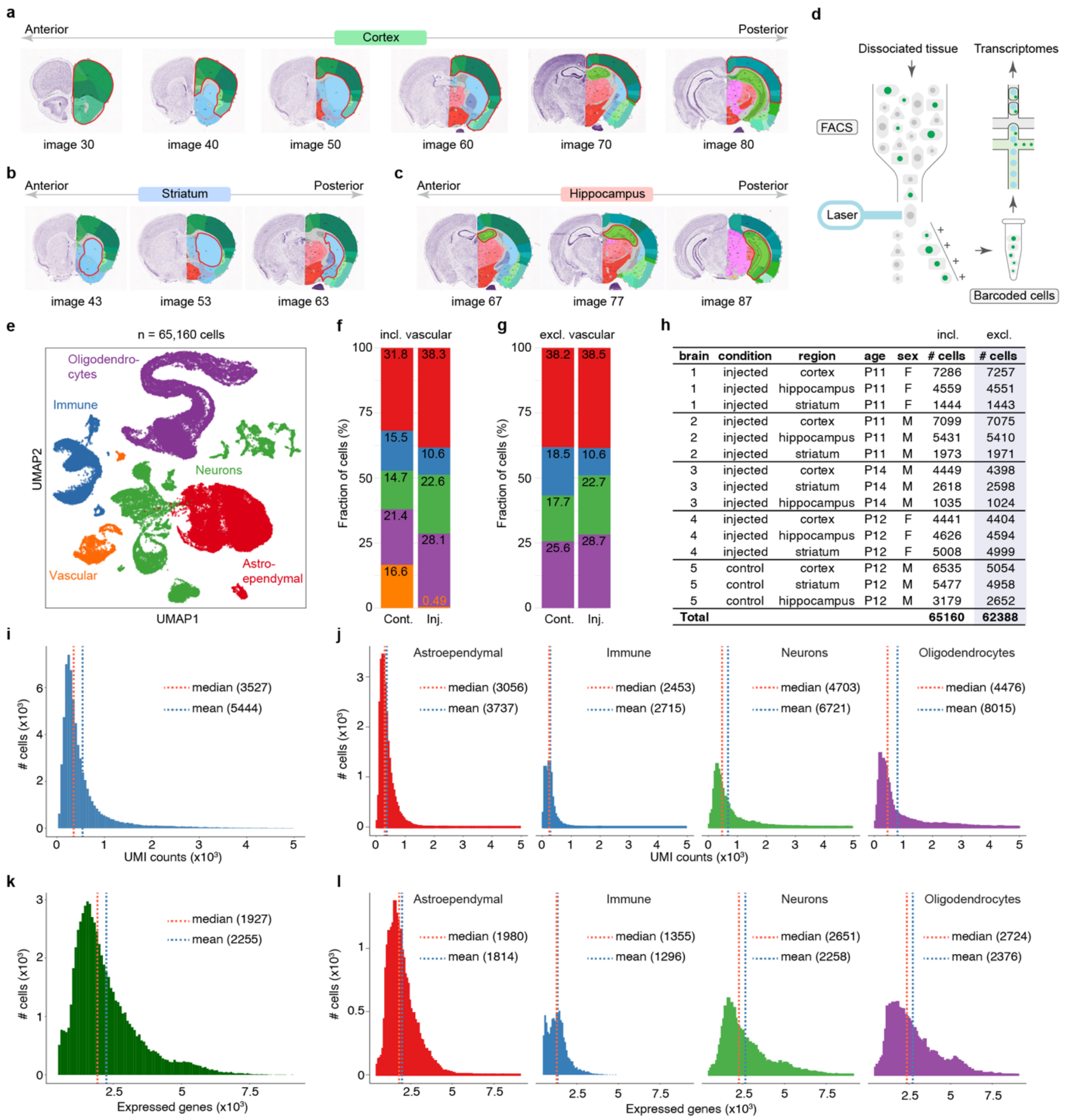


| brain | condition | region | age | sex | incl. # cells | excl. # cells |
|---|---|---|---|---|---|---|
| 1 | injected | cortex | P11 | F | 7286 | 7257 |
| 1 | injected | hippocampus | P11 | F | 4559 | 4551 |
| 1 | injected | striatum | P11 | F | 1444 | 1443 |
| 2 | injected | cortex | P11 | M | 7099 | 7075 |
| 2 | injected | hippocampus | P11 | M | 5431 | 5410 |
| 2 | injected | striatum | P11 | M | 1973 | 1971 |
| 3 | injected | cortex | P14 | M | 4449 | 4398 |
| 3 | injected | striatum | P14 | M | 2618 | 2598 |
| 3 | injected | hippocampus | P14 | M | 1035 | 1024 |
| 4 | injected | cortex | P12 | F | 4441 | 4404 |
| 4 | injected | hippocampus | P12 | F | 4626 | 4594 |
| 4 | injected | striatum | P12 | F | 5008 | 4999 |
| 5 | control | cortex | P12 | M | 6535 | 5054 |
| 5 | control | striatum | P12 | M | 5477 | 4958 |
| 5 | control | hippocampus | P12 | M | 3179 | 2652 |
| **Total** | | | | | **65160** | **62388** |

**Extended Data Fig. 3 | Experimental design, cell numbers and gene expression metrics.**

**a-c,** Each brain was cut using a 1 mm coronal brain slicer and brain regions shown in red contour along the anterior/posterior axis were dissected for tissue dissociation. Image numbers refer to the image number from the Allen Brain Atlas (http://atlas.brain-map.org/atlas?atlas=1#atlas=1). **a**, Cortex samples included the cortical plate (except for hippocampal formation) and most parts of the cortical subplate. **b**, Striatum samples included the caudoputamen and nucleus accumbens. **c**, Hippocampus samples contained the hippocampal formation (except for the entorhinal area). **d**, For each postnatal brain, dissected regions were dissociated separately and barcoded cells were isolated using fluorescence-activated cell sorting (FACS). We used droplet microfluidics (10X Genomics Chromium) to reveal the transcriptomes of barcoded cells. **e**, Initial clustering of cells from both barcoded and non-injected control brains revealed five major cell types annotated and visualized in a UMAP. **f**, **g**, Compared to non-injected control samples (Cont.), barcoded brains (inj.) contained very few vascular cells, because few blood vessels are present at the time of injection (**f**). Therefore, we removed vascular cells from all datasets (**g**). **h**, Table summarizing final cell numbers for each region and replicate. **i, j**, Number of transcripts (unique molecular identifiers, UMIs) per cell for the entire dataset (**i**) and for each major cell type (**j**). **k, l**, Number of expressed genes per cell for the entire dataset (**k**) and for each major cell type (**l**).

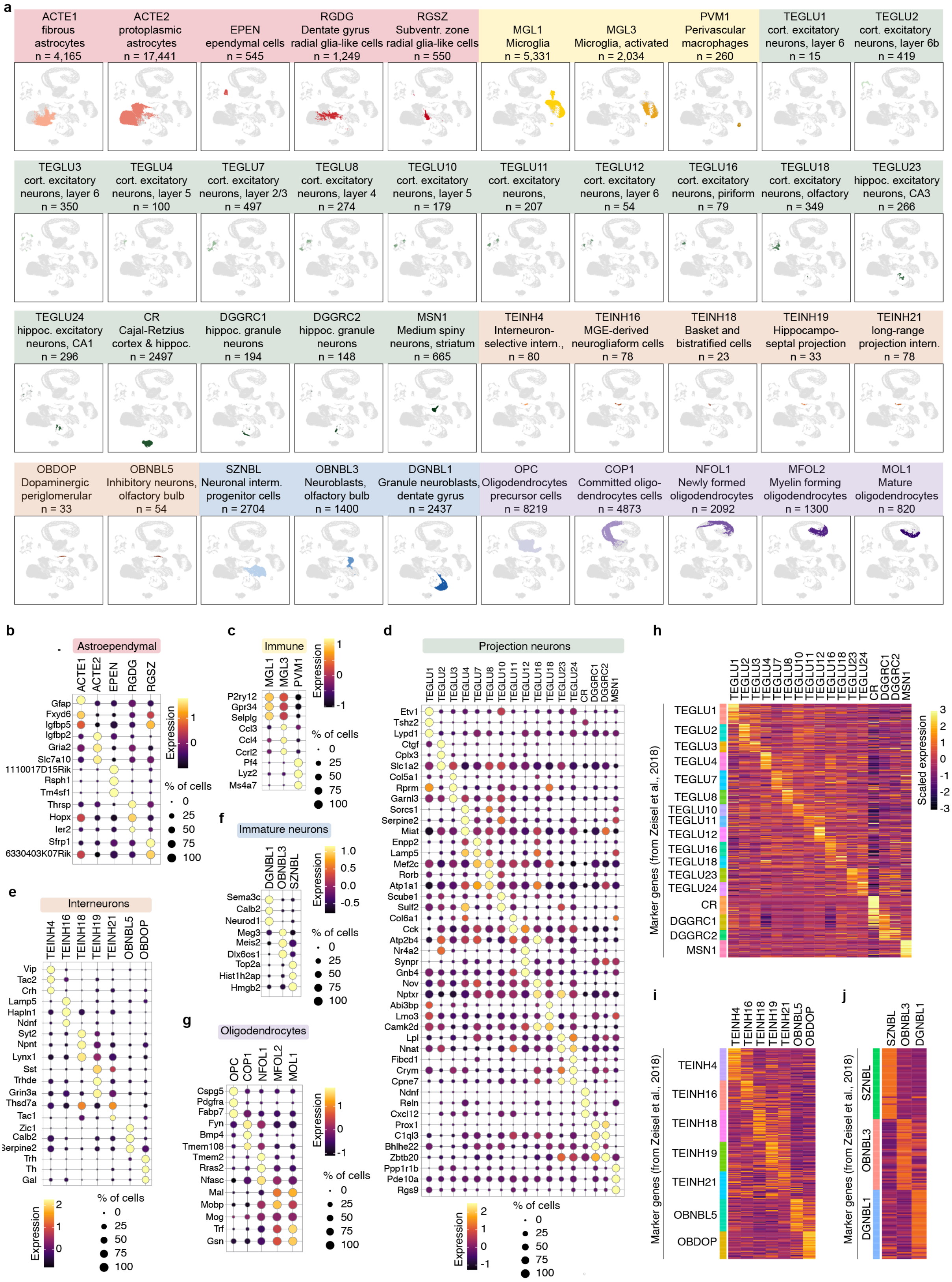


**Extended Data Fig. 4 | Cell types and marker genes.**

**a**, Separate UMAP visualizations for all cell types and corresponding number of cells per type identified in this study. Colors indicate six broader cell type classes: astroependymal (reds), immune (yellows), interneurons (oranges), projection neurons (greens), immature neurons (blues) and oligodendrocytes (purples). **b-g**, Gene expression of markers for each cell type belonging to the six broad cell type classes astroependymal (b), immune (c), projection neurons (d), interneurons (e), immature neurons (f) and oligodendrocytes (g). For each cell type the top three marker genes were identified, and unique genes plotted as dot plots. Expression values represent scaled average gene expression per cell type. **h-j**, We used the same mnemonic identifiers from a previous mouse brain atlas [2] to annotate cell types found in our study. Each heatmap shows the expression for the unique differentially expressed genes (rows) from the mouse brain atlas for the corresponding cluster (columns) from this study for projection neurons (**h**), interneurons (**i**) and immature neurons (**j**).

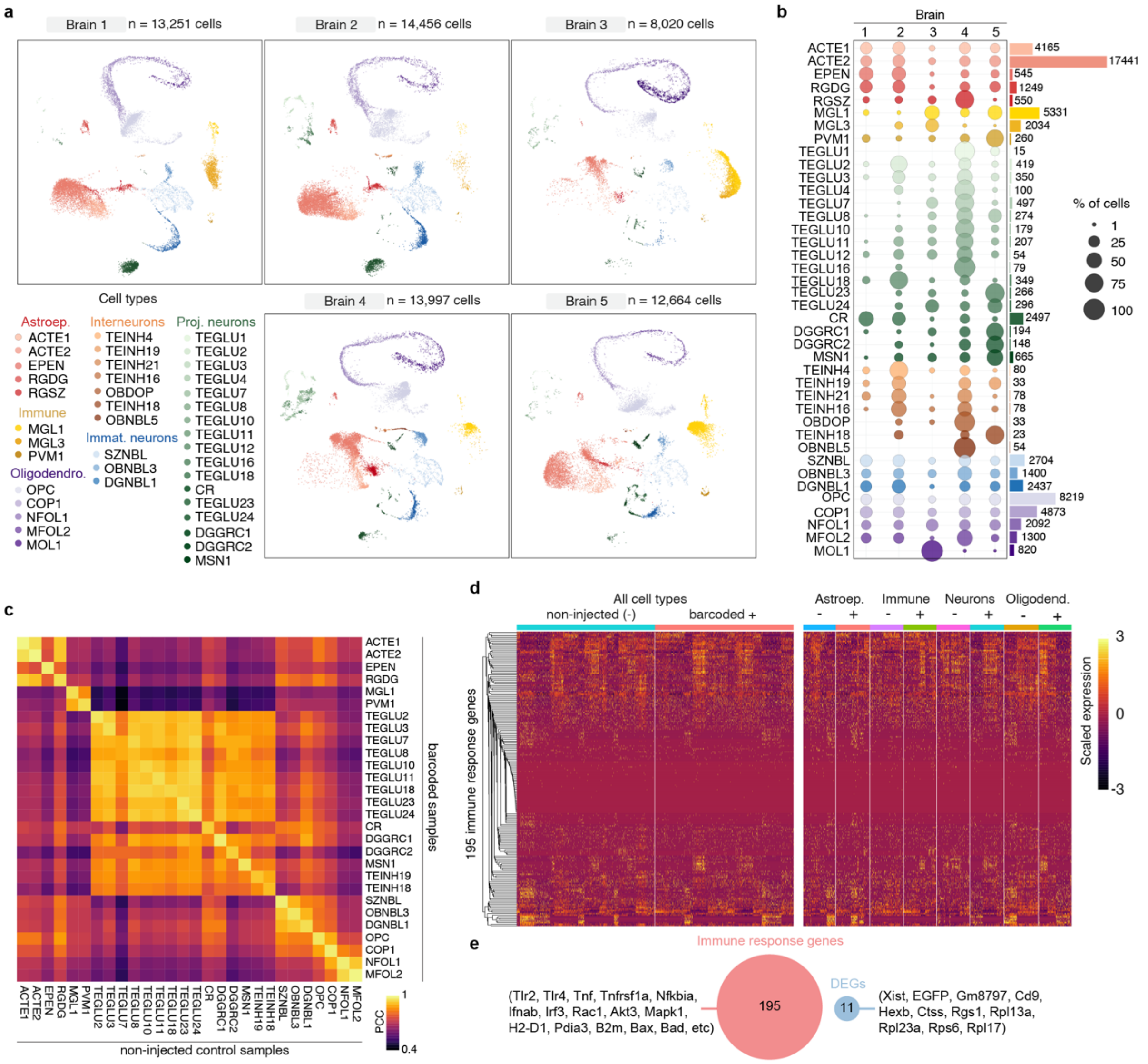


**Extended data Fig. 5 | Lentiviral barcoding does not perturb cell physiology.**

**a,** UMAP visualizations for all 40 cell types split by biological replicate (brain). **b**, Barcoded cells (from brains 1-4) and control cells (from brain 5) are distributed relatively evenly across each cell type. Some variability is expected due to variations in progenitor labeling, cell type sampling and slight age differences between mice used. Dot plot shows fraction of cells per cell type across all brains. Bar plot shows total number of cells per cell type. **c**, Gene expression is similar between control and barcoded cells. Heatmap showing Pearson correlation coefficients (PCC) between average gene expression values for each cell type from barcoded and control brains. A high PCC value indicates similar gene expression patterns between both conditions. Clusters containing at least 5 cells per condition and cell type were analyzed. **d,** Immune response genes are not upregulated in barcoded cells. Heatmap displaying gene expression for 195 immune response genes (KEGG pathway “Human immunodeficiency virus 1 infection”, mmu05170) for an equal number of single cells grouped by condition (left) or grouped by cell type and condition (right). **e**, Only few genes are differentially expressed between barcoded and control cells and do not overlap with immune response genes. Venn diagram showing all differentially expressed genes between barcoded and control cells for each cell type and their non-overlap with immune response genes.

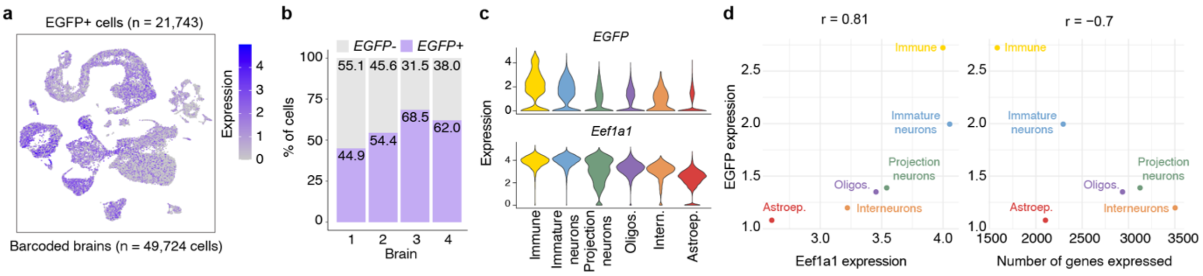


**Extended data Fig. 6 | EGFP expression metrics from synthetic EF1a promoter. a**, UMAP embeddings and normalized EGFP expression levels in all cells (n = 49,724) isolated from postnatal mouse brains that were injected with EF1a-H2B-EGFP-cloneID libraries at E9.5. A total of 21,743 cells contained at least one EGFP transcript. **b**, Bar plots showing the proportion of EGFP transcript positive cells per brain ranging from 44.9% to 68.5% (57.4% ± 10.2%, mean ± SD, n = 4 brains). **c**, Violin plots showing EGFP (top) and Eef1a1 (bottom) expression levels for each broad cell class sorted by decreasing average EGFP expression. **d**, Left, scatter plot showing a high Pearson correlation (r = 0.81) between average expression levels for EGFP and Eef1a1 for each cell class indicating that the synthetic EF1a promoter recapitulates endogenous Eef1a1 expression patterns albeit at lower levels. Right, scatter plot showing a low Pearson correlation (r = –0.7) between EGFP expression levels and average number of genes expressed for each cell class.

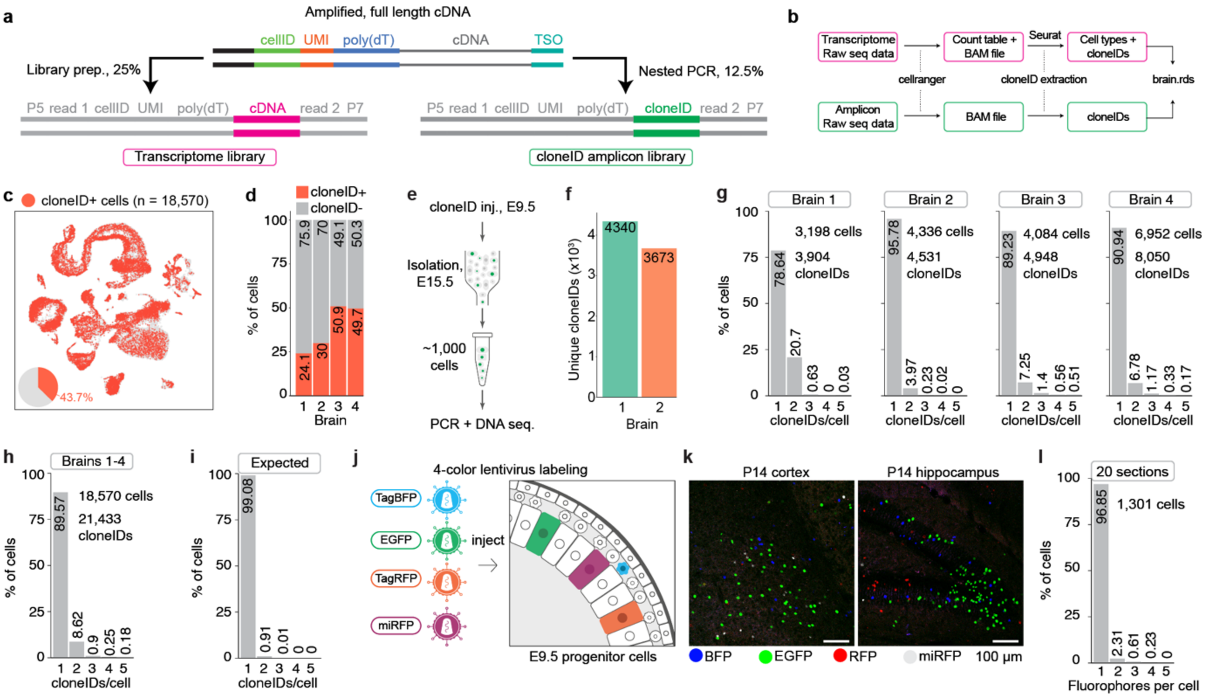


**Extended data Fig. 7 | cloneID sequencing and expression metrics.**

**a**, cloneID sequences are obtained from a transcriptome library and an amplicon library. To enrich for cloneID sequences, we prepared a second library using a targeted, nested PCR approach from full length cDNA obtained during the first steps of transcriptome library preparation. **b**, Simplified workflow for cloneID extraction from amplicon and transcriptome raw sequencing data. We used cellranger (10X Genomics) for raw sequence alignment of both transcriptome and amplicon data and Seurat [3] for analysis of transcriptome data (cell filtering, clustering, etc). We then extracted genetic barcodes (cloneIDs) from aligned sequences (BAM file) corresponding to transcriptome and amplicon data for high quality cells. **c,** UMAP embeddings for all cells (n = 49,724) isolated from barcoded brains and cloneID containining cells highlighted in red (n = 18,570). **d**, Bar plots showing the proportion of cloneID positive cells per brain ranging from 24.1% to 50.9% (38.4% ± 11.8%, mean ± SD, n = 4 brains). **e**, **f,** Targeted DNA sequencing of cloneIDs does not reveal more cloneIDs per cell than expected from scRNA-seq. Bulk DNA sequencing of cloneID sequences from EGFP+ cells isolated six days after lentivirus library injection into the E9.5 mouse brain (c) revealed that the number of cloneIDs ranges from 3,673 to 4,340 unique sequences for 4,000 cells (d). **g, h**, Most cells express only one cloneID upon lentiviral barcoding. Histogram showing the number of cloneIDs per cell for all cloneID positive cells from each brain (g) separately and a summary of all brains (h). **i,** Expected distribution of cloneID copy numbers among transduced cells assuming an idealized model of transduction [4]. Based on the previously observed transduction rate of 1.83%, we expected that >99% of transduced cells contain only cloneID. **j-l**, Most cells express only one fluorophore upon 4-color lentivirus injection. Multicolor labeling of E9.5 progenitor cells with a high titer lentivirus library encoding four different fluorophores (**j**) rarely results in co-expression of more than one fluorophore in the same cell of the postnatal mouse brain (**k, l**) confirming that cloneIDs are quantitatively captured using single cell transcriptomics.

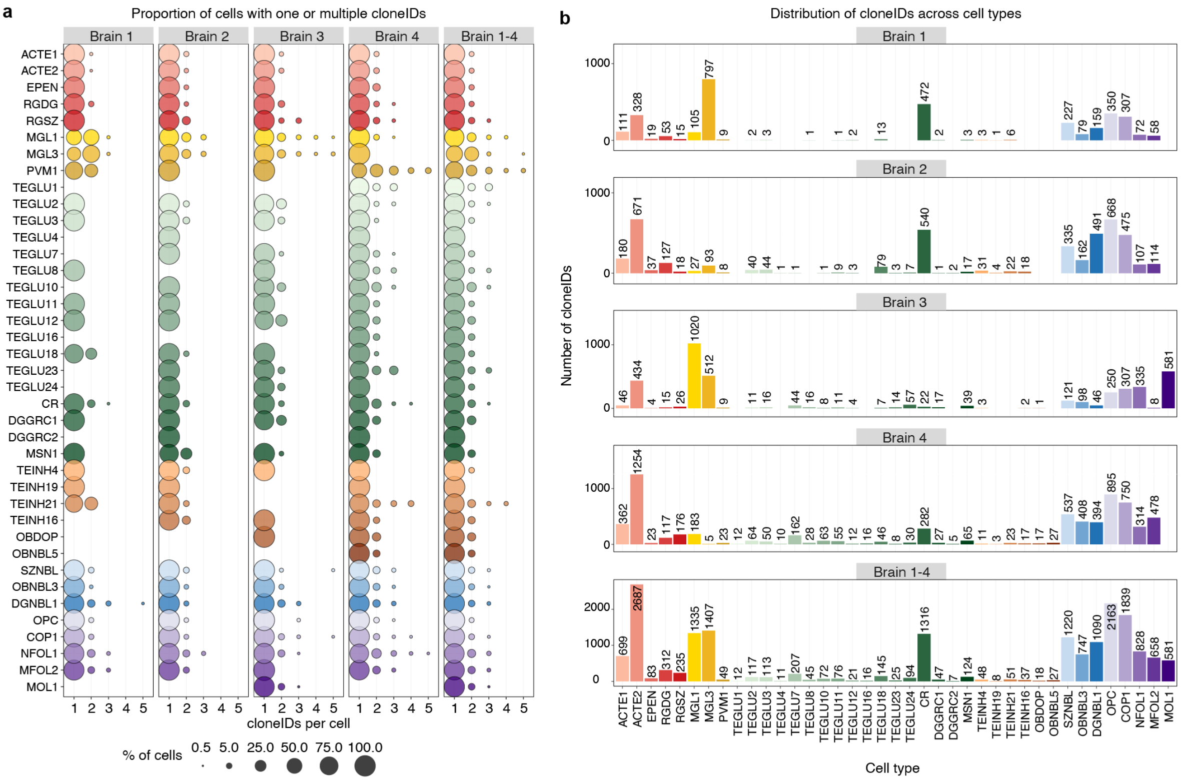


**Extended data Fig. 8 | Distribution of cloneIDs across cell types and brains.**

**a**, Proportion of cells expressing one or multiple cloneIDs. Dot plots show that most cell types express only one cloneID while some cell types express multiple cloneIDs across barcoded brains. **b**, Bar plots showing that the total number of cloneID positive cells varies among cell types and across brains.

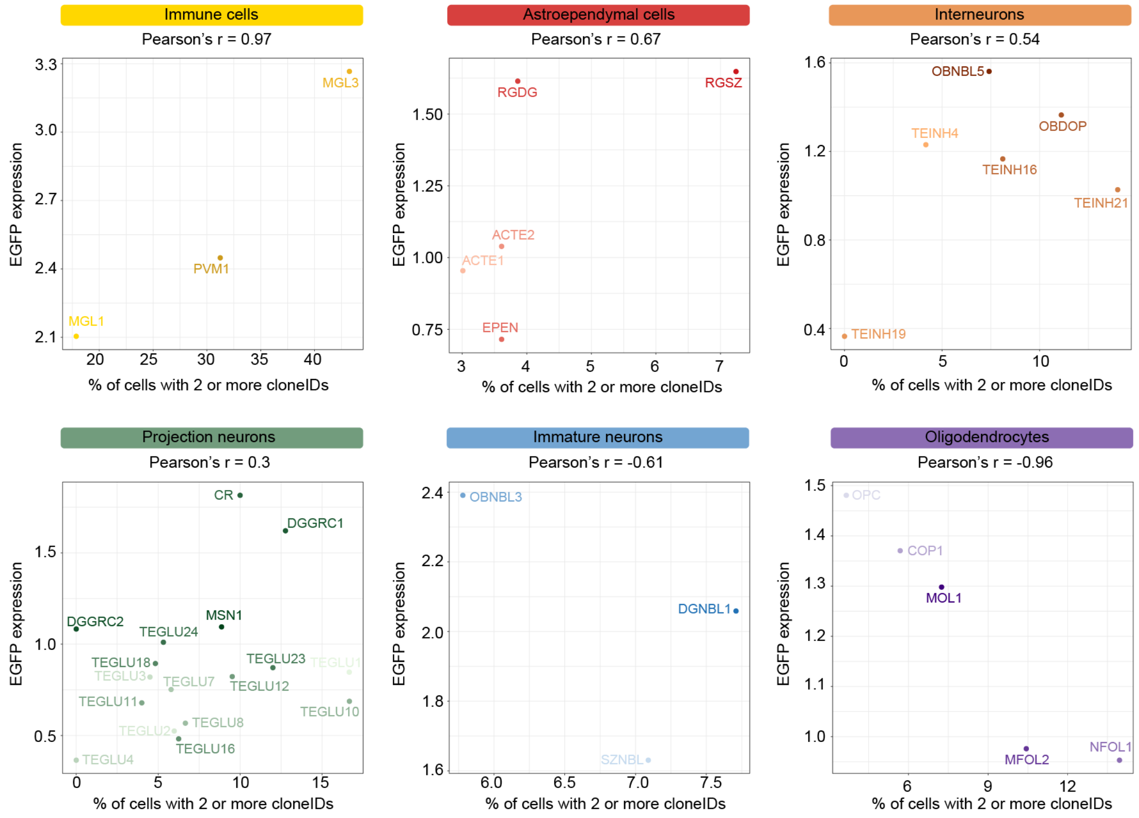


**Extended data Fig. 9 | Correlation between fraction of cells with multiple cloneIDs and cloneID expression levels.**

Scatter plots showing the relation between average EGFP expression level (y-axis) and fraction (%) of cells with 2 or more cloneIDs (x-axis) for each identified cell subtype grouped by major cell class ordered by decreasing Pearson correlation coefficients (r). Correlation coefficients vary greatly depending on cell type and indicate strong positive correlation (immune cells), moderate positive correlation (astroependymal cells, interneurons), weak positive correlation (projection neurons), moderate negative correlation (immature neurons) and strong negative correlation (oligodendrocytes) between EGFP-cloneID expression levels and fraction of cells with 2 or more cloneIDs.

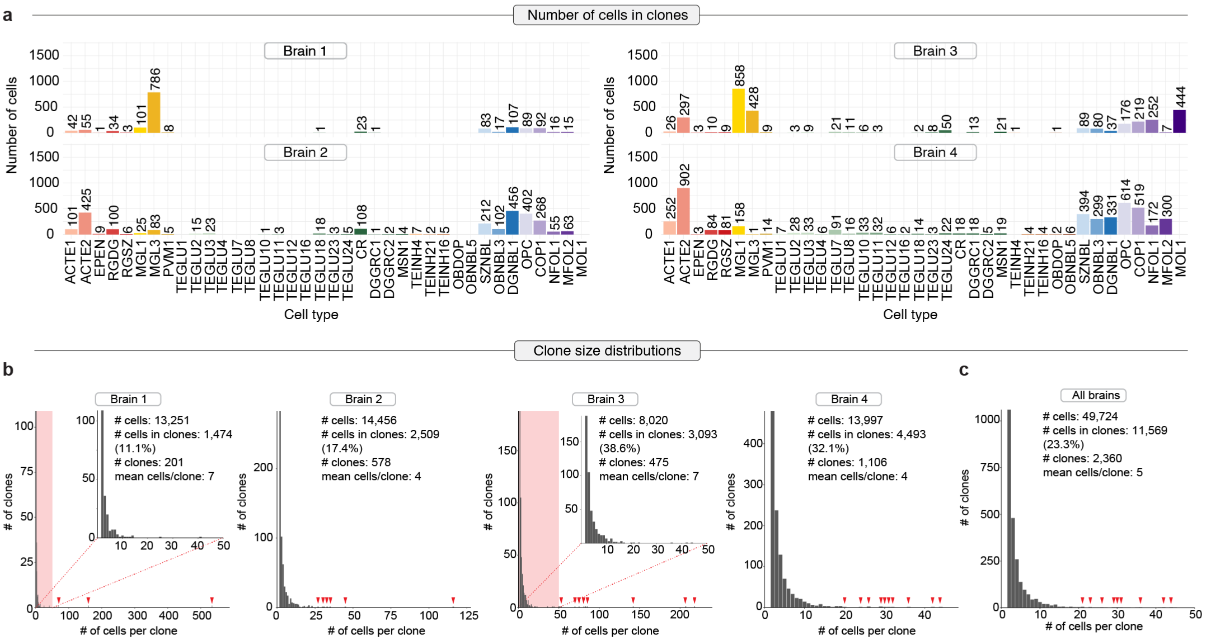


**Extended data Fig. 10 | Clone calling metrics for each brain.**

**a**, Bar plots showing the number of cells contained in clones for cell type and brain. **b**, Histograms and key summary metrics showing the clone size distribution for each brain separately. Shown is the entire distribution of clone sizes for clones containing at least two cells and a magnification (red area) for clone sizes between 2 and 50 cells per clone for brains 1 and 3 that contained a wide range of clone sizes. Red arrow heads indicate the size of rare clones occurring at low frequency. **c**, Histogram and key summary metrics showing the clone size distribution for all clones reconstructed from all brains. Because most clones contained less than 50 less, we displayed only clones ranging from 2 to 50 cells in the histograms.

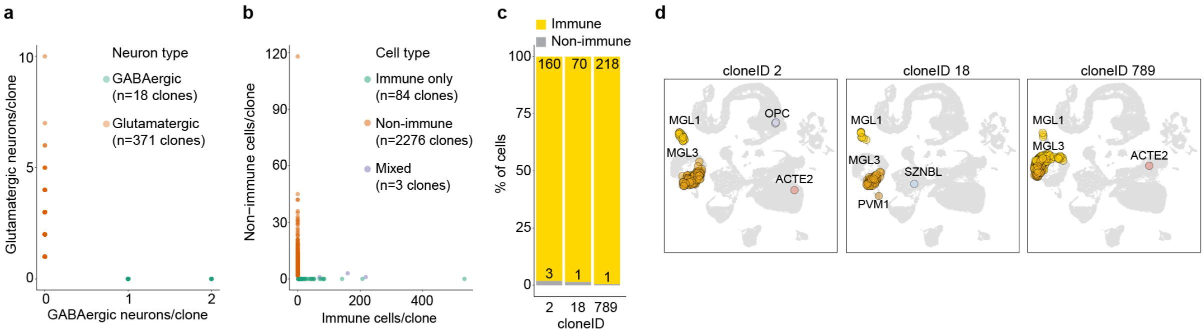


**Extended data Fig. 11 | Cell types of different origin do not share cloneIDs.**

**a**, Scatter plot showing the number of cells in clones containing inhibitory neurons (x-axis, green, n = 18 clones) or excitatory neurons (y-axis, red, n = 371 clones). We never observed a shared cloneID between these two cell types. **b**, Scatter plot showing the number of cells in clones containing mesoderm-derived immune cells (x-axis, green, n = 84 clones) or neuroectoderm-derived non-immune cells (y-axis, red, n = 2,276 clones). We rarely observed mixed clones (n = 3 clones) containing cells of both lineages. **c**, Bar plot showing proportion and total number of cells per type for the 3 mixed clones from (b). Only very large immune clones contain very few non-immune cells indicating a very low rate of contamination. **d**, UMAP visualizations for the 3 mixed clones from (b). The five neuroectoerm-derived cell types observed together with 448 immune cells were two protoplasmic astrocytes (ACTE2), two oligodendrocyte precursor cells (OPC) and one intermediate neuronal progenitor cells (SZNBL).

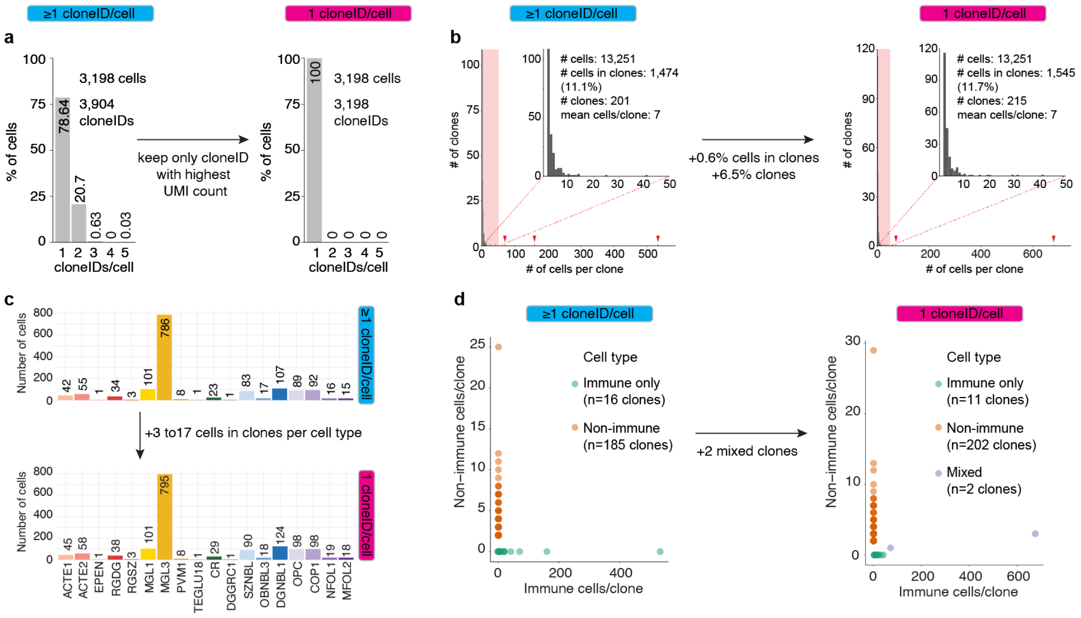


**Extended data Fig. 12 | Removal of multiple cloneIDs from cells leads to higher error in clone reconstruction. a,** When two or more cloneIDs were found in the same barcoded cell of brain 1, we only kept the cloneID with the highest UMI count and removed all other cloneIDs. **b,** Clone size histogram showing that 0.6% more cells are contained in clones and 6.5% more clones are reconstructed when using only a single cloneID. **c,** Depending on cell type, 3 to 17 more cells were found in clones when using only a single cloneID. **d,** Cell types that often express more than one cloneID such as immune cell clones are affected by “lumping” errors leading to less clones with a larger size and a higher number of incorrectly associated neuroectoderm-derived cells.

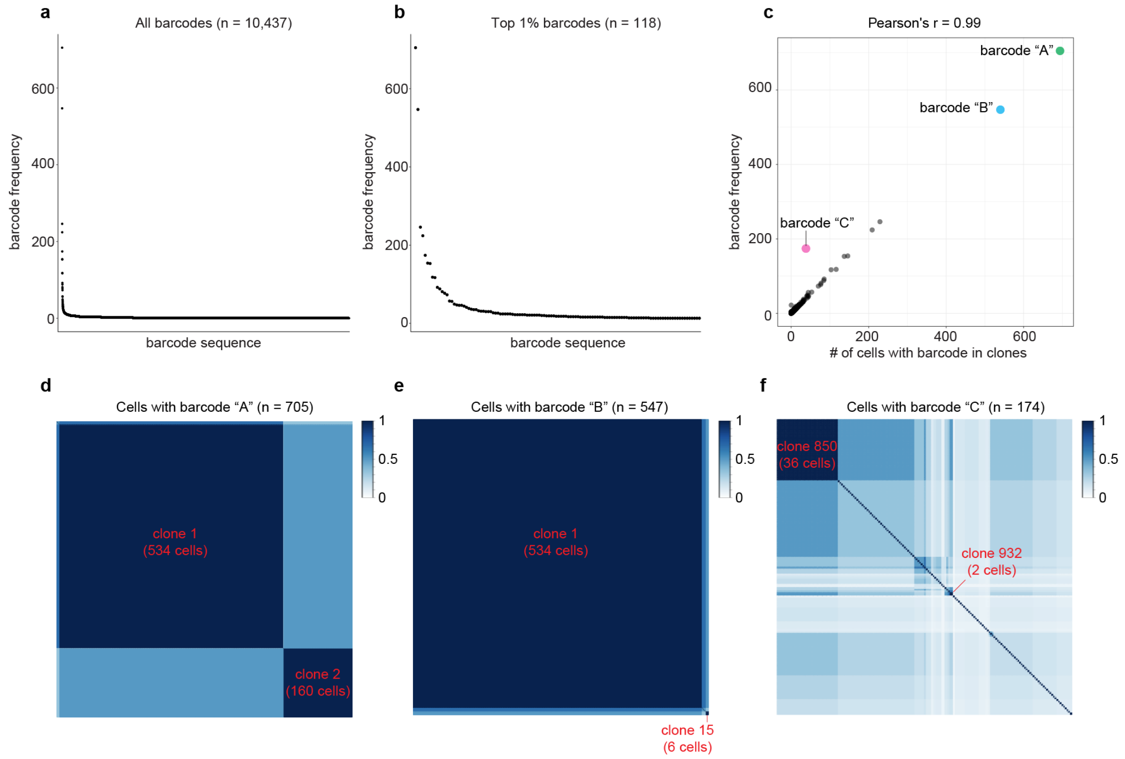


**Extended data Fig. 13 | Frequency distribution of genetic barcodes. a, b,** Rank plots of all unique barcode sequences (a) and the top 1% barcode sequences (b). **c,** A nearly perfect Pearson correlation (r = 0.99) was observed between the number of total occurrences for each barcode sequence (frequency) and the number of cells with a given barcode in clones. Three barcodes named "A", "B" and "C" (nucleotide sequences were omitted for clarity) were highlighted to illustrate the relationships between barcode frequency and number of cells with barcodes in clones in more detail in the panels below. **d-f,** Correlation plots showing Jaccard similarities between all possible pairs of cells expressing three selected barcodes. The two most abundant barcodes "A" and "B" are distributed across three clones (d, e). Clone 1 is defined by the co-expression of both barcodes "A" and "B" and contains 534 cells. Clone 2 expresses only barcode "A" and contains 160 cells. Clone 15 expresses only barcode "B" and contains 6 cells. Barcode "A" is distributed across a total of 705 cells of which the majority (n = 694 cells) is contained in clones while barcode "B" was found in a total of 547 cells of which the majority (n = 540 cells) was detected in clones. The fifth most abundant barcode "C" was found in clone 850 (n = 36 cells) and in clone 932 (n = 2 cells) which co-expresses a second barcode and is therefore considered a separate clone. Out of 174 cells in total expressing barcode "C", only 38 cells are contained in clones. We note that barcode "C" is an outlier barcode since it is the only barcode with a high frequency and low number of cells in clones.

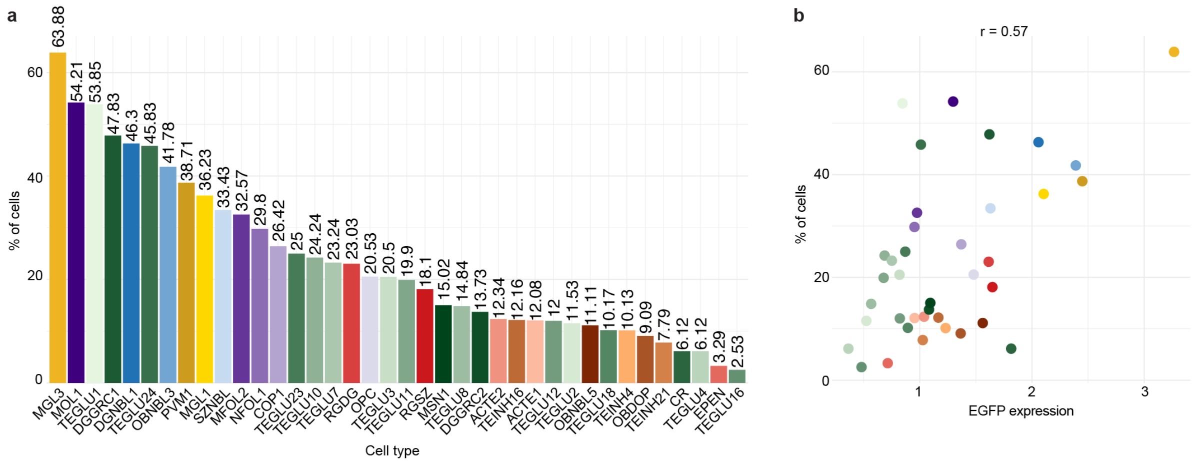


**Extended data Fig. 14 | Number of cells in clones correlates with EGFP expression levels in each cell type.**

**a**, Bar plot showing the proportion of cells in clones for each cell type ordered by decreasing proportion.
**b**, Scatter plot showing a high correlation (Pearson's r = 0.57) between proportion of cells and average normalized EGFP expression for each cell type.

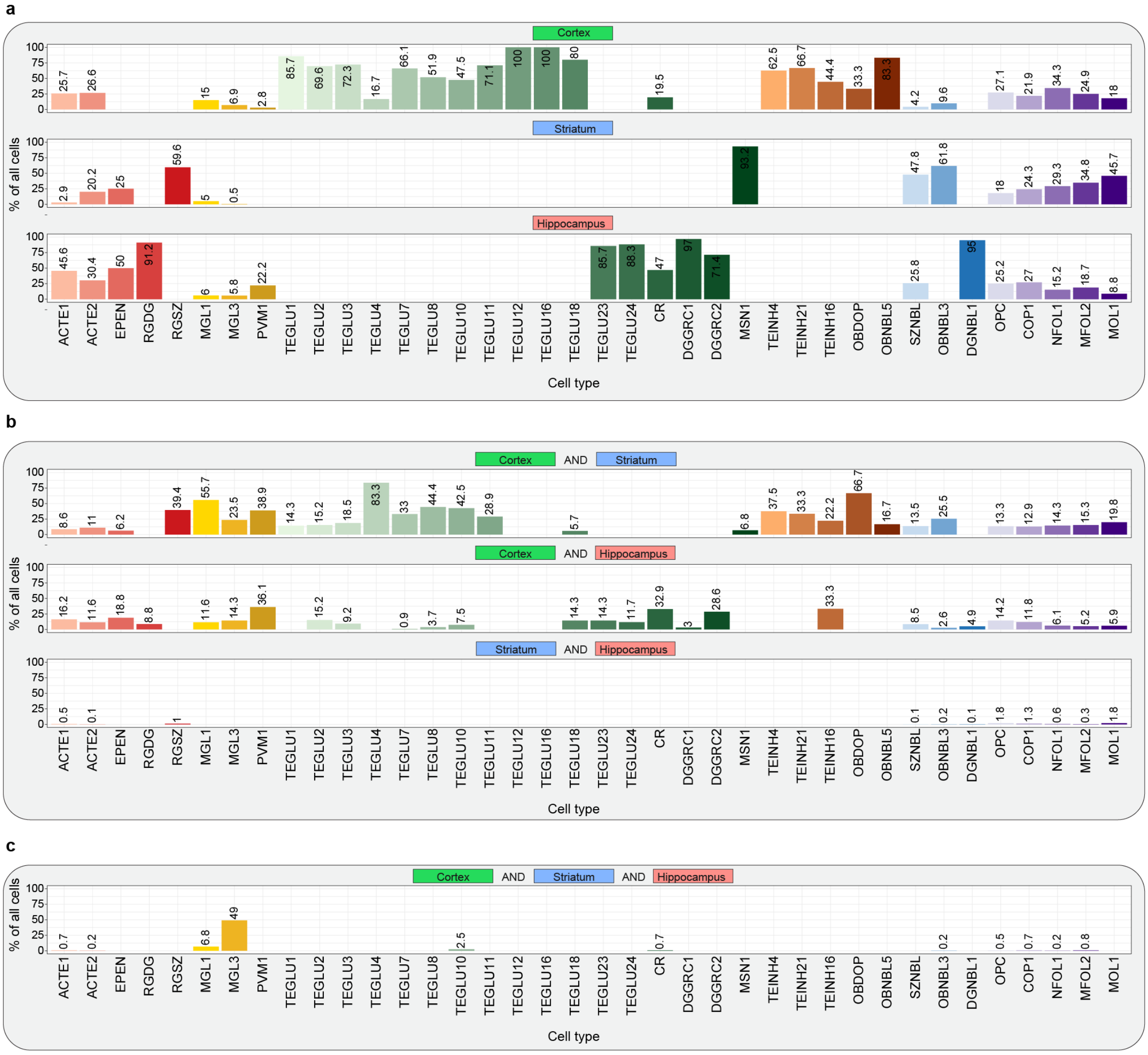


**Extended data Fig. 15 | Proportions of cells per region for each cell type.**

**a-c**, To assess which cell types were associated with dispersed clones and how often they crossed anatomical boundaries between cortex, striatum and hippocampus, we determined the cell type composition of clones located in a single region (a), two regions (b) or all three regions (c).

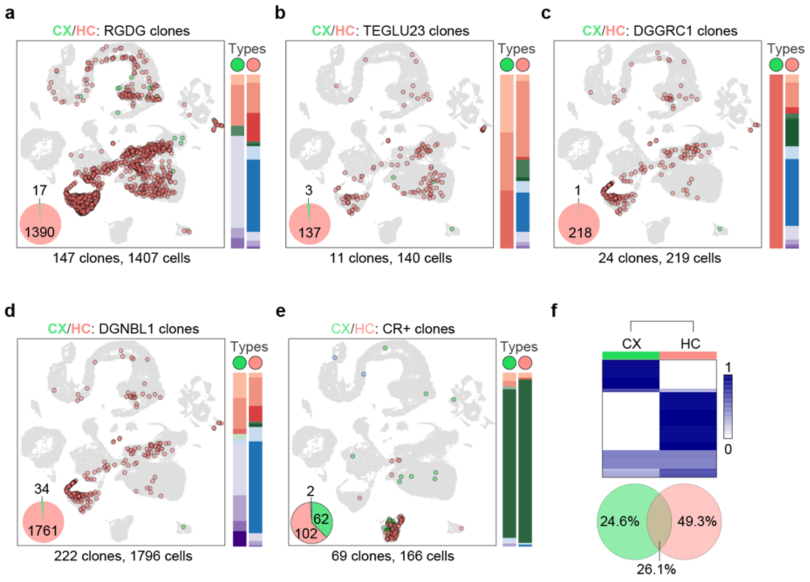


**Extended data Fig. 16 | Regional dispersion of cells that share a cloneID with hippocampus-specific cell types is limited except for Cajal-Retzius cells.**

**a-d,** Most cell types that were specifically found in the hippocampus shared a cloneID with multiple other cell types in hippocampus, but rarely with other cell types in cortex indicating an early segregation of progenitor fields for both regions. Examples for hippocampal cell types are shown for adult neural stem cells in the subgranular zone (a, RGDG), excitatory CA3 neurons (b, TEGLU23), dentate gyrus granule neurons (c, DGGRC1) and granule neuroblasts (d, DGNBL1). **e,** An exception to this pattern are clones with Cajal-Retzius cells (CR). Such clones rarely contained other cell types and often shared a cloneID with CR cells in cortex. **f,** Heatmap showing the proportions of CR cells across both regions for each cloneID. We observed that the 24.6% of cloneIDs accumulated in cortex (CX) only, 49.3% in hippocampus (HC) only and 26.1% are spread across both CX and HC.

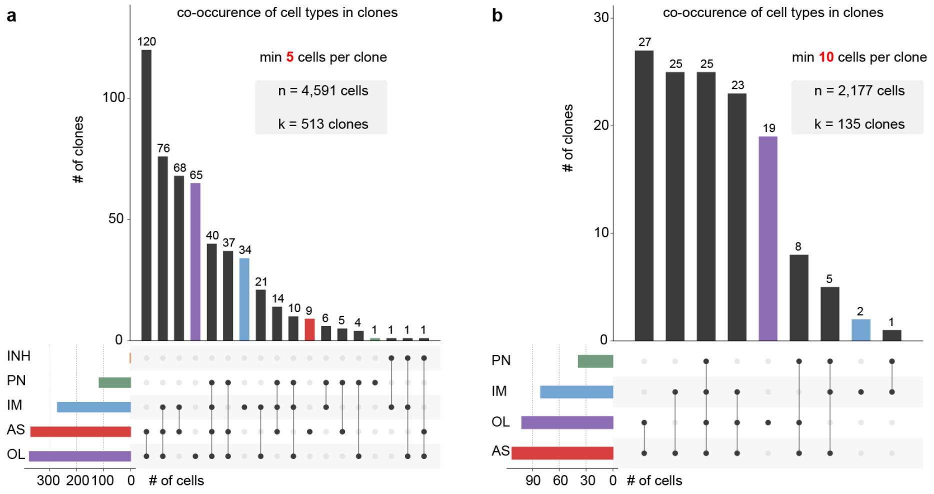


**Extended Data Fig. 17 | Fate distributions of E9.5 neuroectoderm-derived cells.**

**a**, **b**, The number of "minimally uni-potential" clones decreases with increasing clone size. Only clones with neuroectoderm-derived cells containing at least 5 cells per clone (**a**) or at least 10 cells per clone (**b**) are plotted.

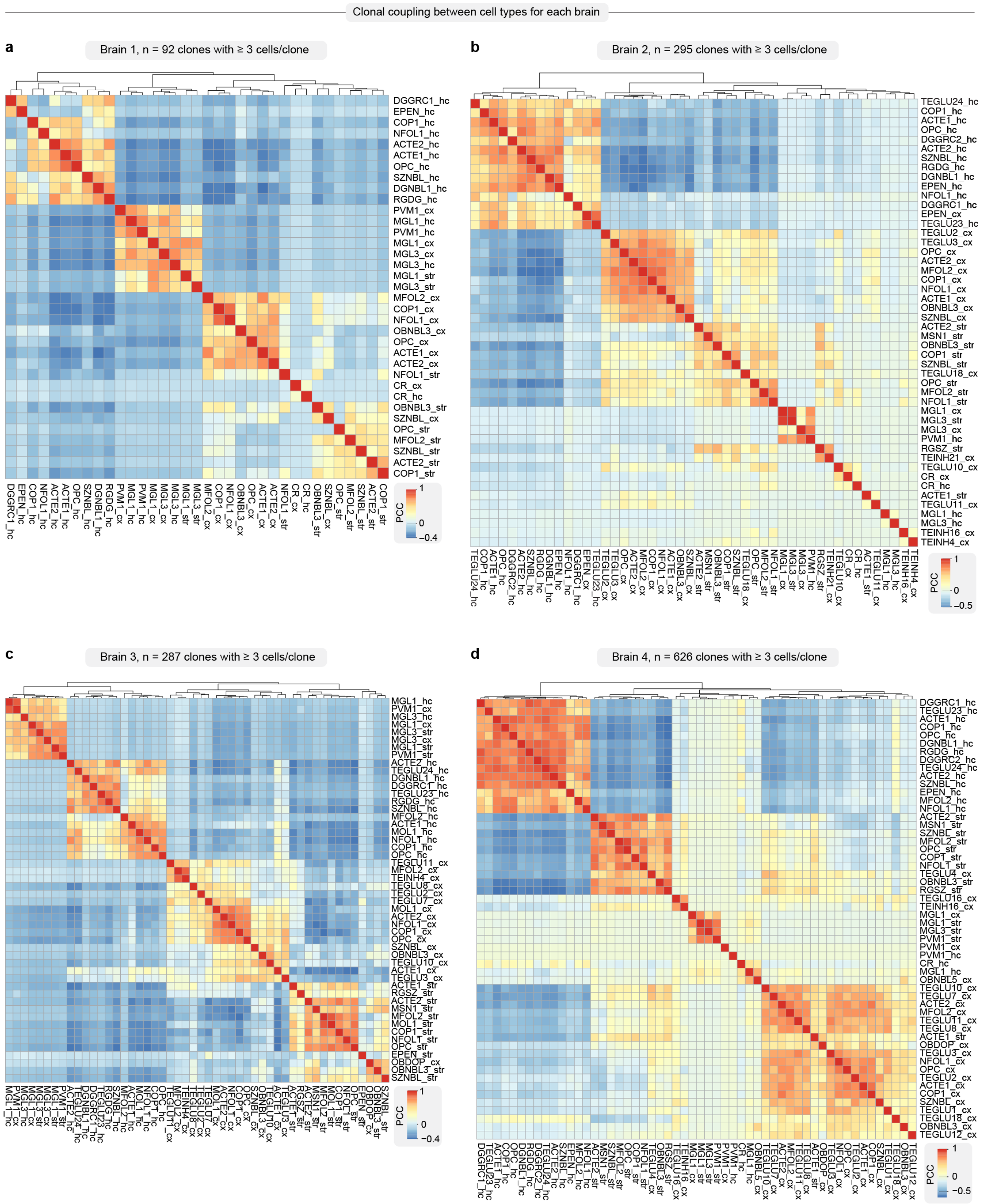


**Extended Data Fig. 18 | Clonal coupling scores for each brain.**

**a-d**, Clonal coupling z-scores defined as the number of shared cloneIDs between all pairs of cell types relative to randomized data were calculated for clonally related cells isolated from brain 1 (**a**), brain 2 (**b**), brain 3 (**c**) and brain 4 (**d**) followed by pairwise correlation of z-scores. Complete-linkage clustering of correlated z-scores revealed structured groups of clonally related cell types as indicated by positive Pearson correlation coefficient (PCC). Clones containing at least 3 cells per clone were considered for each brain.

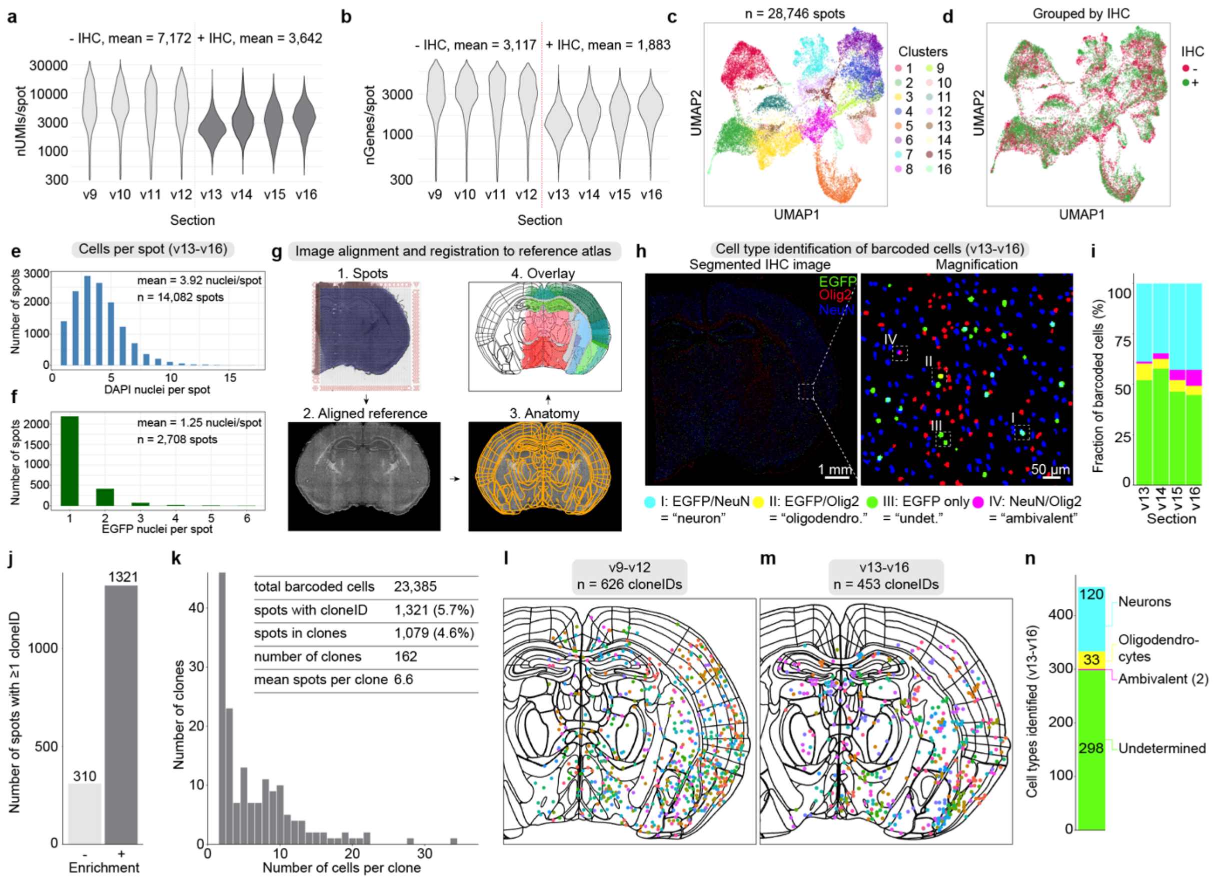


**Extended Data Fig. 19 | Space-TREX enables simultaneous profiling of gene expression, tracking of clonal relationships and phenotyping of cell types *in situ*.**

A total of eight adjacent barcoded brain sections "v9-v16" with a thickness of 10 µm per section were used. Four sections "v9-v12" were processed using the regular spatial transcriptomics workflow and four sections "v13-v16" were used for spatial transcriptomics coupled with cell type identification via immunohistochemical (IHC) staining. **a, b**, The average number of unique molecular identifiers (UMIs) per spot (**a**) as well as the average number of genes detected per spots (**b**) is lower when a section undergoes IHC staining. This is most likely caused by re-folding and re-activation of RNA degrading enzymes in aqueous buffers used for IHC following methanol fixation. **c,** A total of 28,746 spots were sequenced from all eight sections that could be grouped into 16 distinct clusters. **d**, Spots belonging to sections that underwent IHC clustered together with spots from regularly processed sections indicating that IHC staining contains similar molecular information. **e, f,** Histograms showing the number of DAPI+ cells per spot (e) and EGFP+ cells per spot (f) as quantified from immunostained sections v13-v16. **g,** Workflow illustrating image processing for alignment to standardized anatomical reference atlas using WholeBrain[5]. The resulting output contains both the coordinates of spatial transcriptomics spots within the Allen mouse brain reference atlas[6] and each spot is displayed using the regional color code from this reference. **h,** Cell types in EGFP/NeuN/Olig2 triple immunostained tissue sections v13-v16 were identified image segmentation and EGFP+ barcoded cells were classified as "neurons", "oligodendrocytes", "ambivalent" based on the overlap with NeuN, Olig2, NeuN/Olig2, respectively.

Barcoded cells were classified as “undetermined” in absence of overlay with a cell type marker. **i,** **S**ummary of cell types amongst all EGFP+ barcoded cells. **j,** Targeted PCR on full length cDNA from gene expression libraries was used to achieve >4-fold enrichment of cloneIDs resulting in a total of 1,321 spots with ≥1 cloneID. **k,** Distribution of clone sizes for all reconstructed clones. **l, m,** All cloneIDs projected on reference representing non-immunostained (**l**) and immunostained (**m**) sections. **n,** Number of barcoded spots that contained a cell of the type “neuron”, “oligodendrocyte”, “undetermined” or “ambivalent”.

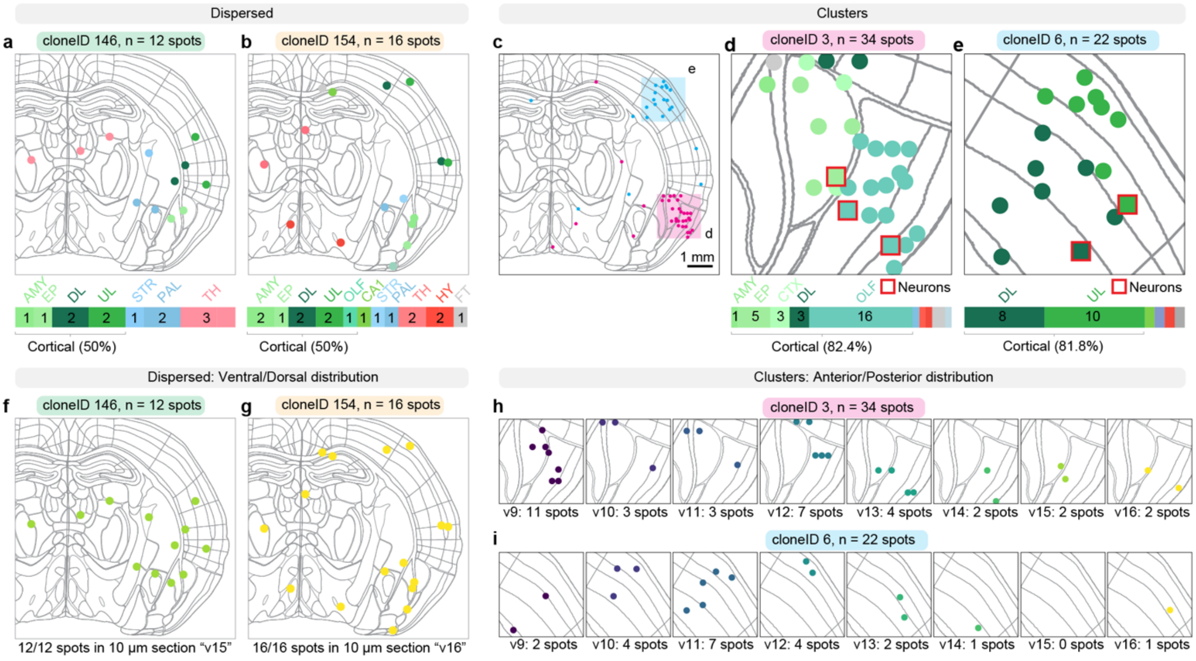


**Extended Data Fig. 20 | Clonal dispersion along dorsoventral and anterior-poserior axis. a, b,** Examples of dispersed clones with detailed regional color code of spots. **c-e**, Examples of clustered clones containing color-coded regional information and the cell type "neuron" encoded as red squares. **f, g,** Cells of dispersed clones spread across the dorsoventral axis within the same 10 µm section of the mouse brain. **h, i,** Cells from clustered clones spread from the most anterior to the most posterior brain section spanning a region of 80 µm.

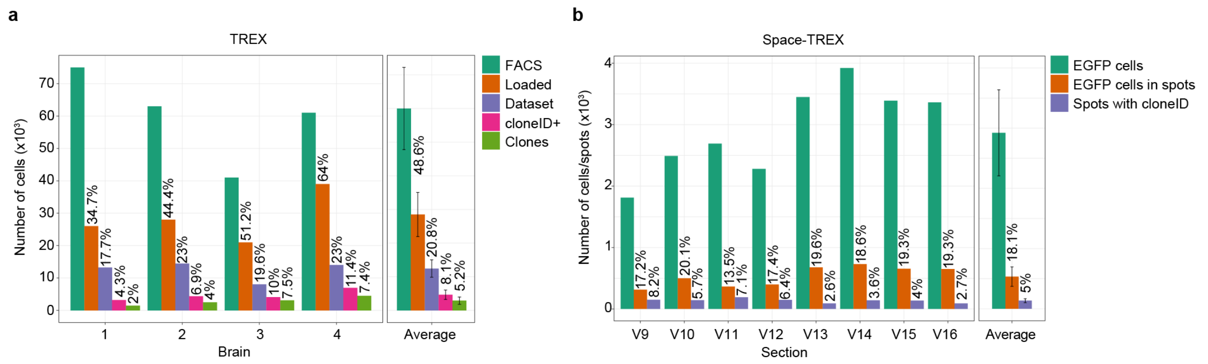


**Extended Data Fig. 21 | Cell loss and barcode dropouts.**

**a,** Bar plots showing the recovery rate of labelled cells and barcodes at each step of the TREX protocol. Upon tissue dissociation isolated all EGFP+ cells contained in the suspension using bulk sorting (darkgreen bar, "FACS") and sorted cells were recovered via centrifugation. All recovered cells were transferred into the wells of a 10X Chromium chip (orange bar, "Loaded") for droplet encapsulation. Following quality control and filtering, a subset of cells was contained in the final dataset (purple bar, "Dataset"). A large fraction of cells had a barcode (magenta bar, "cloneID+") of which many were contained in clones defined as a group of at least two cells sharing the same cloneID (lightgreen bar, "Clones"). The total number of cells is given on the y-axis and each brain as well as the average values for all brains are indicated on the x-axis. The number above each bar represents the total number of cells proportion of cells in that group relative to all cells counted in FACS. **b,** Bar plots showing the recovery rate of barcodes at each step of the Space-TREX protocol. Based on widefield images recorded for each section before library preparation, we could determine the total number of EGFP+ cells in a section (darkgreen bar, "EGFP cells") and the number of those cells located in spots containing capture probes (orange bar, "EGFP cells in spots"). We displayed the final number of spots containing a cloneID (purple bar "Spots with cloneID"). The total number of cells is given on the y-axis and each section as well as the average values for all sections are indicated on the x-axis. The number above each bar represents the proportion of cells/spots in that group relative to all EGFP cells counted.

| | Chan et al., Nature, 2019 | Weinreb et al., Science, 2020 | Pei et al., Cell Stem Cell, 2020 | Bowling et al., Cell, 2020 | He et al., bioRxiv, 2020 | Ratz et al, 2021 (this study) |
|---|---|---|---|---|---|---|
| **Model system** | early embryogenesis | Hematopoietic stem cells (*in vitro* and transplanted) | Hematopoietic stem cells (*in vivo*) | Hematopoietic stem cells (*in vivo*) | Neurogenesis *in vitro* (organoids) | Neurogenesis (*in vivo*) |
| **Species** | Mouse | Mouse | Mouse | Mouse | Human | Mouse |
| **Barcoding approach** | Piggy-bac transposition of multi-copy CRISPR recorder with 3 gRNA targets | Lentiviral integration of random 28 bp barcodes | Cre-driven random deletion or inversion of 9 unique loxP site-flanked DNA blocks located in Rosa26 locus | Dox-inducible CRISPR recorder with 10 gRNA targets spread across Rosa26 and Col1a1 loci | Sleeping beauty transposition of multi-copy CRISPR recorder with 1 gRNA target and 10 bp “integration barcode” | Lentiviral integration of random 30 bp barcodes |
| **Barcode recovery rate** | 15.8-73.7% | 37.7-63.4% | 5.3-18.7% | 32-63% | 17% for lineage barcodes; 51% for integration barcodes | 11.1-38.6% |
| **Cells per clone** | unknown | 8.4-14.9 (mean) | 1.5-4.3 (mean) | 1-123 | 2-801 (range); 11 (mean) | 2-534 (range); 5 (mean) |

**Extended Data Fig. 22 | Recovery rates and clone sizes for combined profiling of lineages and cell types using scRNA-seq in mammalian model systems.** Compared to five other recent studies in mammals, our approach yields comparable barcode recovery rates and numbers of cells per clone and is the only method that has been extensively characterized to meet the needs for studying neurogenesis *in vivo*. Note that we restricted this analysis to recent papers where the relevant information was accessible. Other approaches exist, such as DNA barcoding *in vivo* and *in vitro* using viral libraries of varying complexity[7-12], but mostly their reliance on DNA sequencing for barcode readout and missing information about clone sizes make it challenging to compare those approaches to scRNA-seq studies. Moreover, while commercially available plasmid libraries e.g. CloneTracker XP™ 50M Barcode-3' Library (Cellecta) for delivery of expressed RNA barcodes exist, it is unclear how well barcode complexity is maintained in the final lentivirus preparation, whether virus preparations with titers required for high density *in vivo* barcoding (>$10^9$ TU/ml) can be obtained and how efficient barcodes can be recovered using scRNA-seq assays given that they are expressed at low levels from a truncated version (EFS) of the EF1α promoter[13].

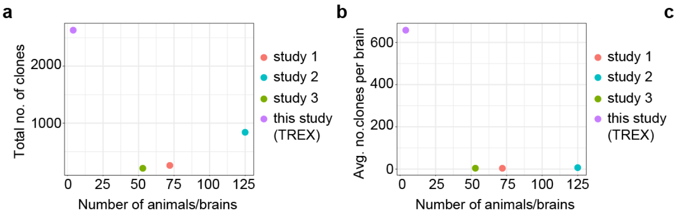


c

| | study 1 | study 2 | study 3 | this study (TREX) |
|---|---|---|---|---|
| **Number of clones** | 260 | 841 | 210 | 2 630 |
| **Number of animals/brains** | 72 | 125 | 53 | 4 |
| **Average number of clones per brain** | 3.61 | 6.73 | 3.96 | 658 |

**Extended Data Fig. 23 | Number of tracked clones using different methods.**
We compared the number of clones detected and animals used between studies that use classical fate mapping methods based on ultra-sparse labeling of progenitors (retroviral tracing, Mosaic Analysis with Double Markers, Cre-induced genetic labeling) and high-density clonal tracing using TREX. **a, b,** Scatter plots showing the total number animals/brains used and the total number of clones detected (a) or the total number animals/brains used and the average number of clones per brain detected (b). **c,** Table summarizing values for all parameters (rows) and studies (columns). The three studies refer to important papers in the field of neurodevelopment published between 2017 and 2020 where the relevant information was accessible: study 1 (ref[14]), study 2 (ref[15]), study 3 (ref[16]).

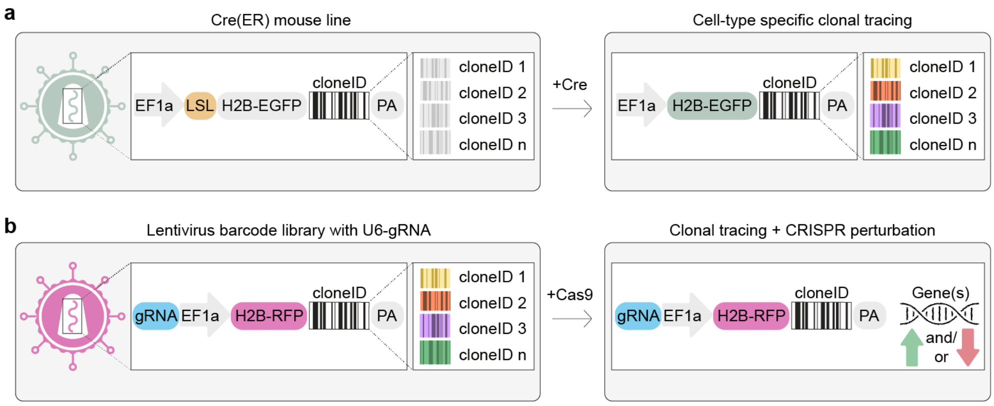


**Extended Data Fig. 24 | Additional lentiviral backbones.**

Next to a backbone enabling constitutive barcode expression in all transduced cell types, we also provide other lentivirus transfer plasmids. **a**, Cell type specific barcoding is enabled by the incorporation of a loxP-stop-loxP (LSL, orange) cassette between EF1a promoter and H2B-EGFP-cloneID transgene. The LSL cassette is removed in presence of Cre recombinase to restrict barcoding to genetically defined, Cre-expressing (precursor) cells. Note that many resources of Cre driver lines for genetic targeting of specific cell types are available[17,18]. **b**, A construct containing a H2B-RFP-cloneID transgene (red) and a U6-gRNA scaffold expression cassette (blue) allows to combine clonal tracing with CRISPR-based gene perturbations upon injection into a Cas9-EGFP mouse[19]. Note that CRISPR can be employed for both target gene upregulation (green arrow) as well as downregulation (red arrow) and multiple genes can be targeted which will allow to couple genetic screens to clonal tracing[20].
